# AT2-intrinsic Z-AAT expression drives conserved inflammatory and proteotoxic stress responses and predisposes to emphysema

**DOI:** 10.64898/2026.04.06.715869

**Authors:** Carly Merritt, Rose P. Griffin, Kristine M. Abo, Joseph E. Kaserman, Pushpinder Singh Bawa, Carlos Villacorta Martin, Feiya Wang, Michael Morley, Michael Cho, Maria C. Basil, Maor Sauler, Andrew A. Wilson

**Affiliations:** Center for Regenerative Medicine of Boston University and Boston Medical Center, Boston, MA; The Pulmonary Center and Department of Medicine, Boston University School of Medicine, Boston, MA; Department of Medicine, Perelman School of Medicine, University of Pennsylvania, Philadelphia, PA; Penn-CHOP Lung Biology Institute, Perelman School of Medicine, University of Pennsylvania, Philadelphia, PA; Channing Division of Network Medicine, Brigham and Women’s Hospital, Harvard Medical School, Boston, MA; Pulmonary, Critical Care, and Sleep Medicine, Yale School of Medicine, New Haven, CT

## Abstract

Individuals homozygous for the *SERPINA1* “Z” mutation with alpha-1 antitrypsin deficiency (AATD) are highly susceptible to emphysema. This predisposition has classically been attributed to a relative deficiency of circulating alpha-1 antitrypsin (AAT) reaching the lungs and associated protease-antiprotease imbalance. Accumulating evidence suggests that the presence of misfolded Z-AAT protein either in the circulation, the lung interstitium, or within resident lung cells could contribute to emphysema pathogenesis. We have shown that type 2 alveolar epithelial cells (AT2s), progenitor cells of the lung alveolus, heterogeneously retain Z-AAT and exhibit a transcriptional disease signature in AATD patient samples. However, a lack of model systems that faithfully recapitulate AT2 biology and associated Z-AAT expression has limited our ability to study this phenomenon. Here, we apply syngeneic induced pluripotent stem cell-derived AT2s (iAT2s) and a novel mouse model featuring AT2-specific inducible human *SERPINA1* expression to interrogate the cell-instrinsic consequences of Z-AAT expression, validating findings in an independent dataset of human COPD lung tissue comparing ZZ to MM *SERPINA1* genotypes. We find further evidence of Z-AAT retention within AT2s and identify shared AT2 transcriptomic disease signatures conserved across model systems, characterized by innate immune and inflammatory signaling, NF-κB activation, and endoplasmic reticulum stress. Mice with AT2-specific Z-AAT expression additionally demonstrate increased susceptibility to elastase-induced emphysema, providing functional evidence for AT2-intrinsic contributions to AATD-associated lung disease. Within iAT2s, a subpopulation of Z-AAT expressing cells exhibits activation of the PERK-eIF2α signaling axis and markers of an alveolar basal intermediate (ABI) state, emerging cell-autonomously in the absence of mesenchymal co-culture.Together, these data provide evidence that Z-AAT expression in AT2s induces heterogenous cell-intrinsic stress responses including proteotoxic stress, inflammatory signaling, and aberrant cell fate adoption, and is sufficient to result in predisposition to injury, supporting a potential contribution of AT2-intrinsic Z-AAT toxicity to human AATD-associated emphysema pathogenesis.

## Introduction

Alpha-1 antitrypsin deficiency (AATD) is caused by mutations in *SERPINA1*, the gene encoding alpha-1 antitrypsin (AAT). AAT, secreted primarily by hepatocytes, is an abundant anti-inflammatory serum glycoprotein that acts as an antiprotease, inhibiting neutrophil elastase in the lungs. AAT is also produced by other immune and epithelial lineages, including neutrophils^1^, alveolar macrophages^2^, monocytes^3^, and airway epithelial cells^4^. The most prevalent disease-causing *SERPINA1* variant, known as the “Z” mutation, is a point mutation that results in an amino acid substitution (G342K) which predisposes Z-AAT proteins to misfold and polymerize within the endoplasmic reticulum (ER)^5,6^. Patients homozygous for the Z mutation (ZZ-AATD) are predisposed to developing both lung and liver disease. While AATD-associated liver disease results as a direct consequence of accumulated Z-AAT polymers within hepatocytes^5,6^, AATD-associated emphysema has been primarily attributed to loss of antiprotease function in the lung and resultant unopposed neutrophil elastase activity leading to pulmonary parenchymal destruction^5,7^.

Several lines of evidence, however, suggest that mechanisms beyond simple loss of function toxicity could contribute to lung disease in AATD. Many severely deficient AATD patients treated with protein replacement “augmentation” therapy continue to experience lung function decline in excess of that associated with normal aging^8, 9^. Z-AAT polymers, including those in the lung, are known to increase activation and trafficking of immune cells^10–12^. In addition, gain-of-function toxicity associated with Z-AAT production has been observed in multiple cell types outside of the liver including circulating monocytes and neutrophils^1, 3^. Recently, we and others have identified a transcriptomic disease signature among resident lung macrophages and type 2 alveolar epithelial cells (“AT2s”) that express mutant *SERPINA1*^13,14^. Because AT2s are facultative progenitors of the lung alveolus, the structure primarily injured in AATD-associated emphysema, establishing the pattern of AAT expression within these cells and delineating any associated consequences of Z-AAT expression is of vital importance in furthering understanding of AATD-associated emphysema pathogenesis.

Model systems commonly applied for studies of AATD pathogenesis are not well-suited to address questions of Z-AAT-associated toxicity in AT2s. Existing mouse models, for example, either lack AAT expression altogether^15^ or lack expression from AT2s^16^. Immortalized cell lines, meanwhile, fail to faithfully recapitulate essential AT2 biological programs^17,18^ or capture patient-to-patient genetic heterogeneity that could underlie commonly observed variation in disease manifestations^19,20^. We and others have applied iPSC-derived AT2s (iAT2s) to model monogenic diseases^21–24^ and functional significance of genes identified through genome-wide association studies^25,26^. Here, building on our prior transcriptional characterization of Z-AAT expressing AT2s in primary human lung tissue, we apply syngeneic ZZ and wild-type (MM) iPSCs to interrogate the cell-instrinsic consequences of Z-AAT expression in human AT2s. To determine these effects *in vivo*, we developed a novel conditional mouse model allowing AT2-specific inducible expression of wild-type or mutant human *SERPINA1*. We validate transcriptional findings from both experimental systems in an independent dataset of single nucleus RNA sequencing from human COPD lung tissue comparing ZZ to MM *SERPINA1* genotypes. Through integrated analysis across all three systems, we identify a conserved transcriptional signature characterized by innate immune and inflammatory signaling and activation of the PERK-eIF2a axis. Finally, we demonstrate that AT2-intrinsic Z-AAT expression is sufficient to increase susceptibility to experimental emphysema *in vivo*, providing functional evidence for a contribution of AT2-instrinsic Z-AAT toxicity to AATD-associated emphysema pathogenesis.

## Results

### Generation of ZZ and MM human iPSCs

We sought to develop an experimental platform that would allow us to determine whether dysregulated processes previously observed in ZZ AT2s from end-stage COPD primary lung tissue (“ZZ-COPD”) might originate early in the disease process or result only later as a consequence of longstanding injury^13^. To do so, we turned to the human iPSC platform that we have previously applied to model disease mechanisms in ZZ liver hepatocytes^27^. Building on an existing repository of iPSCs from ZZ and control (MM) donors^28^, we utilized CRISPR/Cas9 gene editing to either repair or introduce the Z mutation, resulting in syngeneic pairs of ZZ and MM iPSCs from four genetic donors (PiZZ1, PiZZ2, PiZZ6, and BU3, Figure 1A). All resulting lines underwent characterization to confirm successful editing and retention of a normal karyotype (Figure S1A,B).

**Figure 1:**
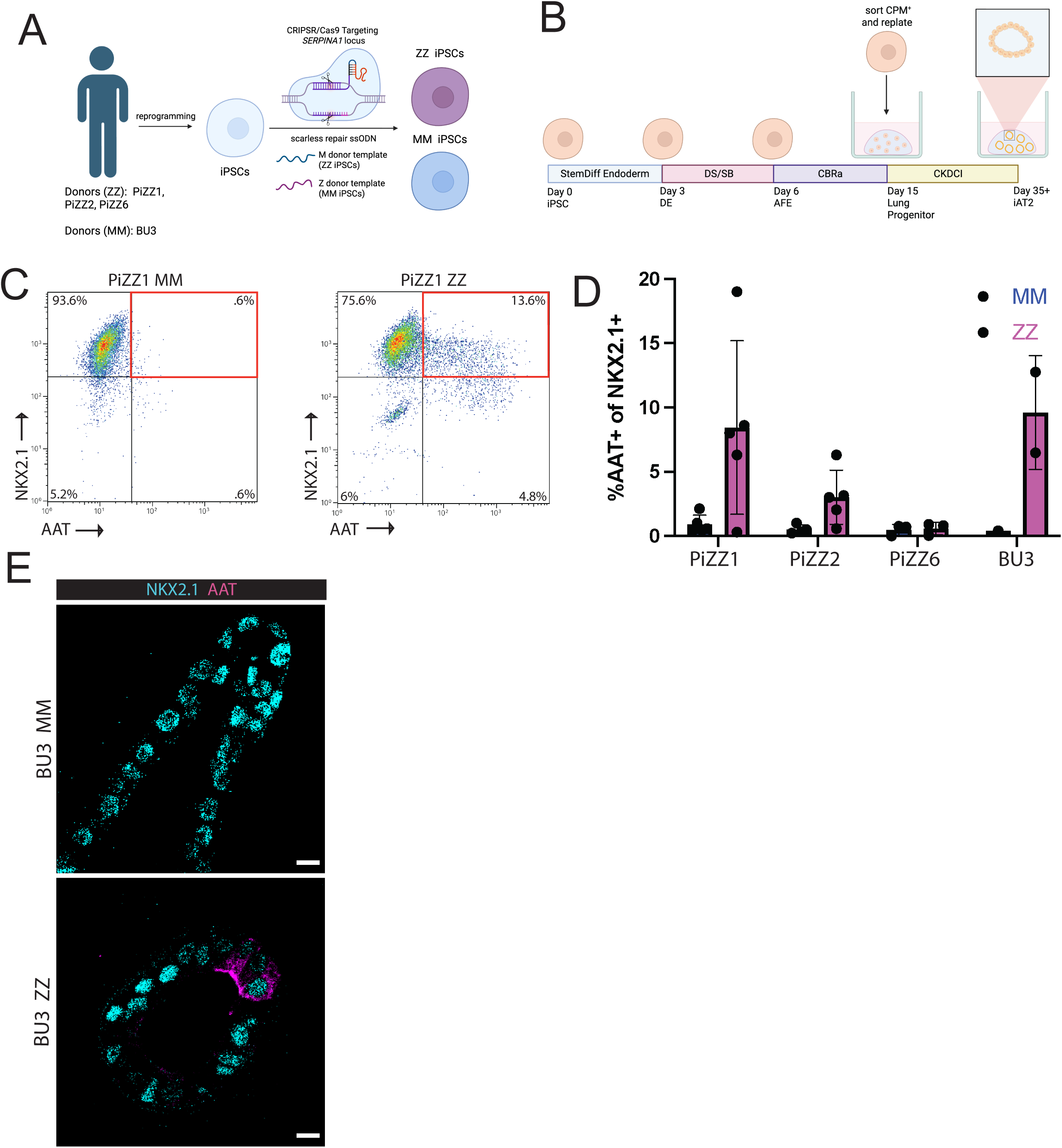
Patient-derived ZZ iAT2s exhibit heterogeneous intracellular AAT retention. A) Targeting strategy for the SERPINA1 locus B) Schematic of directed differentiation protocol for generating iAT2 C) Representative flow cytometry plots of fixed, permeabilized ZZ and iAT2s. D) Mean percent of NKX2.1+ iAT2s staining positive for intracellular A protein in ZZ and MM iAT2s. Each dot represents the mean over one independent differentiation (PiZZ1 n=5, PiZZ2 n=5, PiZZ6 n=3, BU3 n=2). Data represented as mean +/- SD. E) Immunostaining of ZZ and MM iAT2s for AAT and NKX2.1. Scale bar 10 um.

### ZZ iAT2s exhibit increased AAT retention

We have previously identified aberrant retention of AAT protein in a subset of AT2s from ZZ-COPD but not MM-COPD or control lung tissue samples^13^. Because we and others have previously found that iPSC-derived AT2s (“iAT2s”) recapitulate in vivo human biology^23,29–31^, we applied a directed differentiation protocol which has been shown to efficiently generate AT2-like cells that transcriptomically and functionally resemble primary AT2s (Figure 1B, Figure S1C)^21^. We generated iAT2s from the four syngeneic pairs of ZZ and MM lines and stained cells with antibodies to identify intracellular AAT protein in cells expressing the lung-defining transcription factor NKX2.1, quantified by flow cytometry. We repeated this experiment between 2-5 times for each syngeneic pair for a total of 32 independent differentiations (Figure 1D, Figure S1D).

Consistent with our observations in primary lung tissue, we found heterogeneous intracellular AAT protein retention among ZZ iAT2s compared to little or undetectable amounts in MM controls. The proportion of ZZ iAT2s that retained intracellular AAT varied both by differentiation for each individual iPSC line and between iPSC lines (0-19%) derived from different donors. To confirm AAT protein is expressed by epithelial cells of lung rather than non-lung origin using an orthogonal approach, we performed immunostaining for AAT and NKX2.1 on cross sections cut from fixed, embedded iAT2 spheres (Figure 1E, Figure S1E) and again observed positive AAT staining in a subset of NKX2.1^+^ ZZ iAT2s but none in MM iAT2s. Together, these data demonstrate that a subset of ZZ iAT2s aberrantly retain AAT protein intracellularly compared to syngeneic MM controls.

### ZZ iAT2s exhibit heterogeneous transcriptomic evidence of ER stress and inflammation

Having established a patient-specific iAT2 disease model that recapitulates findings from primary human lung samples, we next sought to identify mechanisms and pathways that might be dysregulated by Z-AAT expression. Specifically, we were interested in potential heterogeneity within ZZ iAT2s, as our previous data demonstrated heterogeneous AAT protein retention in both iAT2s and explant lung AT2s in addition to ZZ hepatocytes^13,27^. Using scRNA-seq, we profiled the transcriptome across ZZ and MM iAT2s from the PiZZ1 line (Figure 2A). UMAP clustering showed cells clustering by genotype (Figure 2B). Differential gene expression demonstrated 2360 upregulated genes and 1524 downregulated genes in ZZ compared to MM iAT2s (Figure S2A) Louvain clustering via Seurat identified 5 clusters, 2 composed of MM and 3 composed of ZZ iAT2s (Figure 2C). We analyzed canonical AT2 markers and found similar expression across clusters except cluster 4, which expressed lower levels of AT2 marker genes. No expression of gastric, liver, or lung marker genes associated with non-AT2 epithelial lineages were identified across clusters (Figure 2D). To identify the drivers of clustering, we visualized the top 50 DEGs for each cluster combined with Hallmark FGSEA (Figure 2E,F). Based on these analyses, we annotated the clusters as 0: MM homeostatic, 1: ZZ IFN signaling, 2: MM proliferative, 3: ZZ proliferative, and 4: ZZ proteotoxic stress. Both the MM and ZZ proliferative clusters differentially expressed genes associated with cell cycle (*MKI67, TOP2A)* and were enriched for the Hallmark gene sets “E2F Targets” and “G2M checkpoint” (Figure 2E-G). All three ZZ clusters exhibited increased relative expression of IFN-associated genes (*IFI44L, OAS2, IFI6)*, and cluster 1 was specifically enriched for the Hallmark gene sets “IFN Alpha Response” and “IFN Gamma Response” (Figure 2E-G). Distinguishing it from the other ZZ clusters, cluster 4 specifically demonstrated increased expression of ER chaperone genes (*HSPA6, HSPA1B)* and the PERK-UPR genes *DDIT3*, *GADD45B,* and *ATF3* and was enriched for Hallmark pathways “TNFα Signaling via NF-κB” and “Hypoxia” (Figure 2E-G). qRT-PCR of an independent differentiation confirmed increased expression of key ER stress genes in ZZ compared to MM iAT2s, including *PPP1R15A, DDIT3, GADD45A, ATF6,* and *HERPUD1* (Figure S2C-G). Overarching differences between ZZ and MM iAT2s were reflected by enrichment of Hallmark gene sets such as “IFN alpha signaling”, “IFN gamma signaling”, “Inflammatory Response” and “TNFα signaling via NF-κB” (Figure S2B) consistent with findings observed in primary AT2s^13^ suggesting dysregulation of innate immune and inflammatory signaling in Z-AAT expressing cells.

**Figure 2:**
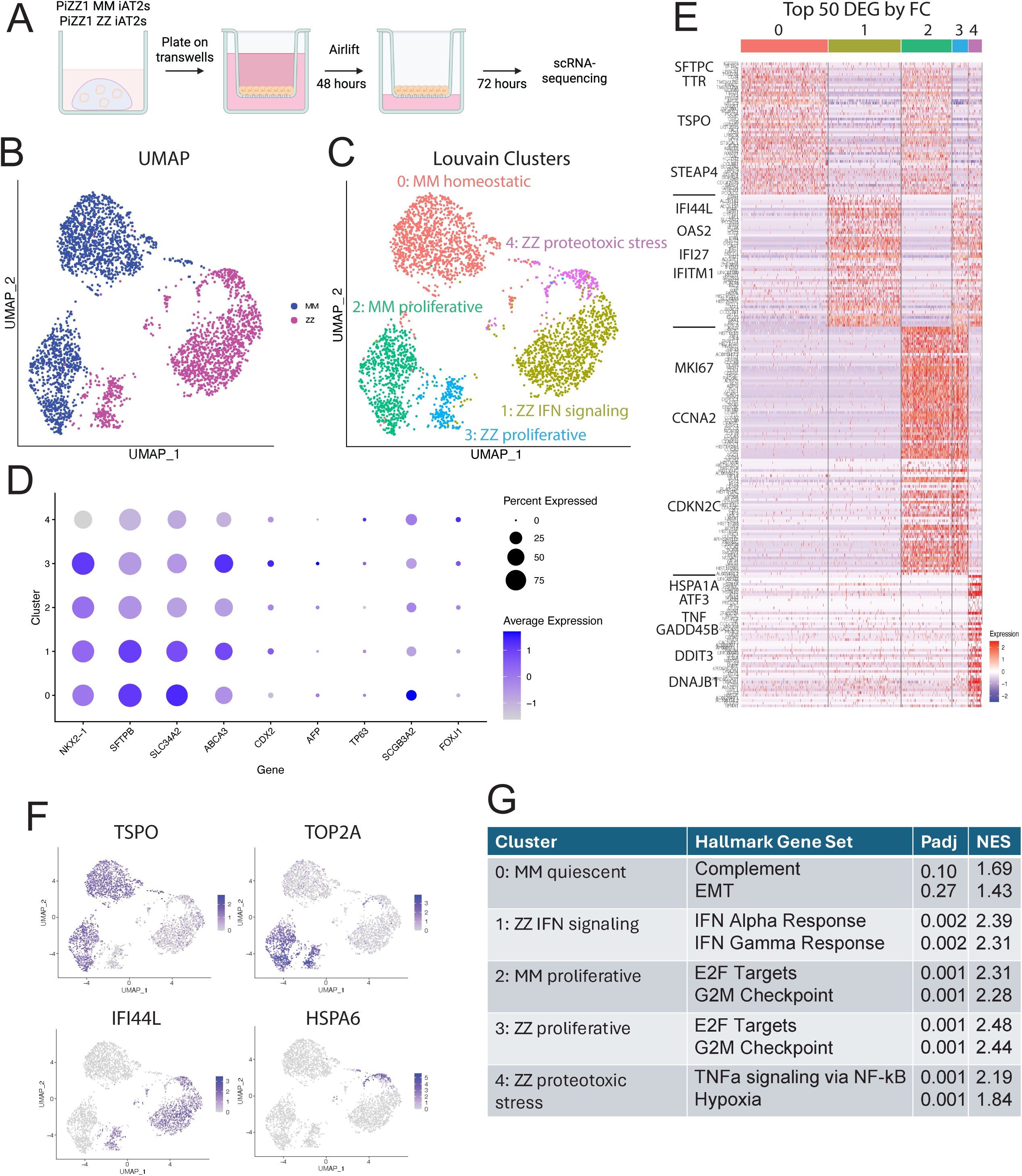
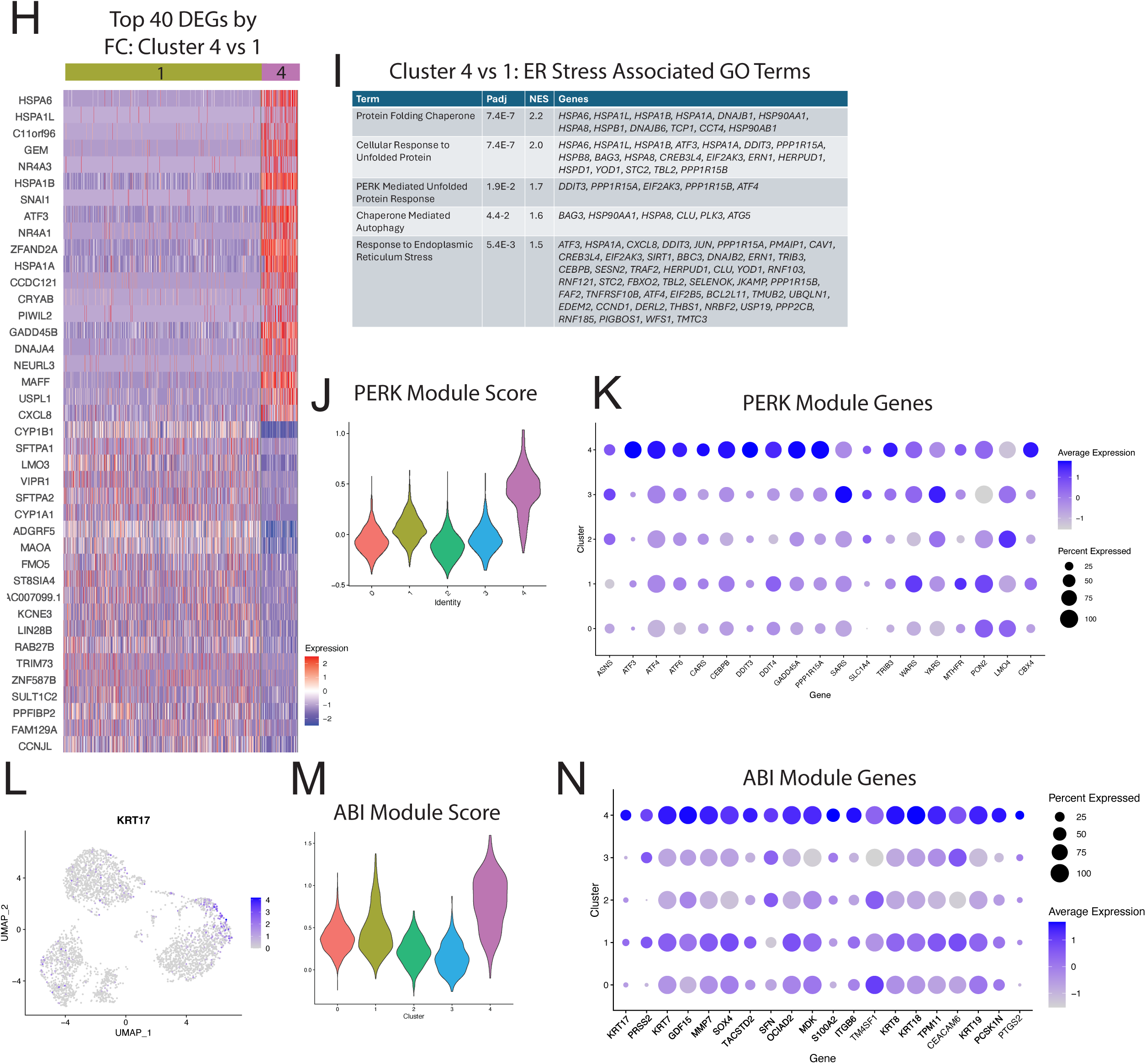
scRNA-seq demonstrates a heterogeneous transcriptional cellular stress response among ZZ iAT2s. A) Experimental plan. PiZZ1 ZZ and MM iAT2s were plated on transwells, airlifted, and underwent scRNA-sequencing. B) Projection of PiZZ1 ZZ and PiZZ1 MM iAT2 scRNA-seq transcriptomes using Uniform Manifold Approximation Projection (UMAP). C) UMAP by Louvain clustering identifies 5 clusters. D) Average expression and frequencies of select lung, AT2, intestinal, hepatic, and airway markers across the 5 clusters. E) Top 50 DEGs per cluster as ranked by log2FC. A subset of upregulated genes is highlighted with large font. F) UMAP of select cluster-specific associated DEGs. G) Top two Hallmark gene sets by each cluster as ranked by NES using GSEA of all DEGs. H) Top 20 DEGs as ranked by log2FC directly comparing clusters 1 and 4. I) Top ER stress associated GO terms for cluster 4 compared with cluster 1 by GSEA. J) Violin plot of PERK module score across all clusters. K) Dot plot projection for PERK branch-specific genes. Differentially expressed genes are bolded (padj < .05, log2FC > .25). L) Feature plot of KRT17 in UMAP space. M) Violin plot of ABI module score across clusters. N) Dot plot projection for ABI-specific genes. Differentially expressed genes are bolded (padj < .05, log2FC > .25). A-N) PiZZ1 line, n=1

To further explore heterogeneity within the ZZ iAT2s, we directly compared the gene expression pattern between clusters 4 (“proteotoxic stress”) and 1 (“IFN signaling”). We first examined the top 40 DEGs between clusters, and found the proteotoxic stress cluster again expressed higher levels of ER chaperone genes *(HSPA6, HSPA1B, HSPA1A),* stress-responsive genes (*ATF3, GADD45B*), and the inflammatory cytokine *CXCL8* while the IFN signaling cluster demonstrated higher expression of genes associated with AT2 identity (*SFTPA1, SFTPA2*) and xenobiotic metabolism (*CYP1A1, CYP1B1*) (Figure 2G). To determine whether Z-AAT-associated proteotoxicity activated ER stress pathways in iAT2s, we next performed Gene Ontology enrichment analysis and found cluster 4 to be enriched for ER-stress associated GO terms “Protein Folding Chaperone”, “Cellular Response to Unfolded Protein”, “PERK mediated Unfolded Protein Response”, “Chaperone Mediated Autophagy”, and “Response to Endoplasmic Reticulum Stress” (Figure 2I). Having previously observed specific activation of PERK in a subset of iPSC-hepatocytes^27^, we were interested to determine whether a similar phenomenon might occur in iAT2s. We applied a gene module for the PERK arm of the UPR, and in fact found a significantly increased PERK module score^32^ in the ZZ proteotoxic stress cluster compared to all other clusters (Figure 2J). To identify drivers of this module score we looked at expression of the individual genes composing the module and found them to be broadly upregulated in cluster 4 compared to all other clusters (Figure 2K). Gene modules for ATF6 and IRE/XBP1-associated signaling did not reveal enriched expression of transcripts representative of other UPR arms (Figure S2H,I). Further examination of differential gene expression between clusters 4 and 1 revealed significant upregulation of keratin family members including *KRT17* that, together with other markers, has been found to identify an aberrant epithelial cell state in the injured distal lung, variably referred to as transitional, alveolar basal intermediates (ABI)^33^, aberrant basaloid^34^, KRT5-/KRT17+^35^, and other terms. Examining the expression of genes upregulated in KRT5-/KRT17+ lung epithelial cells from patients with pulmonary fibrosis, we observed significant enrichment in cluster 4 (Figure 2L-N). Together, these data demonstrate a Z-AAT-associated transcriptional disease signature among iAT2s with heterogeneous downstream consequences in a subset of Z-AAT expressing cells that include expression of proteotoxic stress genes, activation of the PERK-eIF2⍺ signaling axis, and the emergence of an ABI state.

### A novel mouse model of AT2-specific human AAT expression

To determine the consequences of Z-AAT expression in AT2s in vivo, we developed a novel mouse model in which human *SERPINA1* cDNA (either wild-type M or mutant Z) and eYFP conjugated to a self-cleaving 2A peptide are expressed downstream of a floxed stop codon in the *Rosa* locus (*Rosa26*^LSL-*SERPINA*1–2A-eYFP^). Prior literature has largely applied two primary murine models for studies of AATD. These include the PiZ transgenic mouse, which expresses Z-AAT in hepatocytes and macrophages but not lung epithelium^16^ and the *Serpina1a-e-KO* mouse, which lacks expression of endogenous *Serpina1* following deletion of a conserved sequence from five murine paralogs^15^. While the *Serpina1* knockout mouse does develop emphysema, neither of these mice expresses Z-AAT in lung epithelium. We crossed *Rosa26*^LSL-*SERPINA*1–2A-eYFP^ mice with SFTPC^Cre/ERT2^ mice to generate offspring which express human *SERPINA1* and eYFP specifically in AT2s upon tamoxifen-induced recombination (hereafter referred to as AT2^SERPINA1-ZZ^ and AT2^SERPINA1-MM^) (Figure 3A). While these mice retain expression of murine *Serpina1* and thus do not fully recapitulate AATD patient gene expression patterns, they allowed us to isolate the effects of Z-AAT expression in the absence of systemic AAT deficiency.

**Figure 3:**
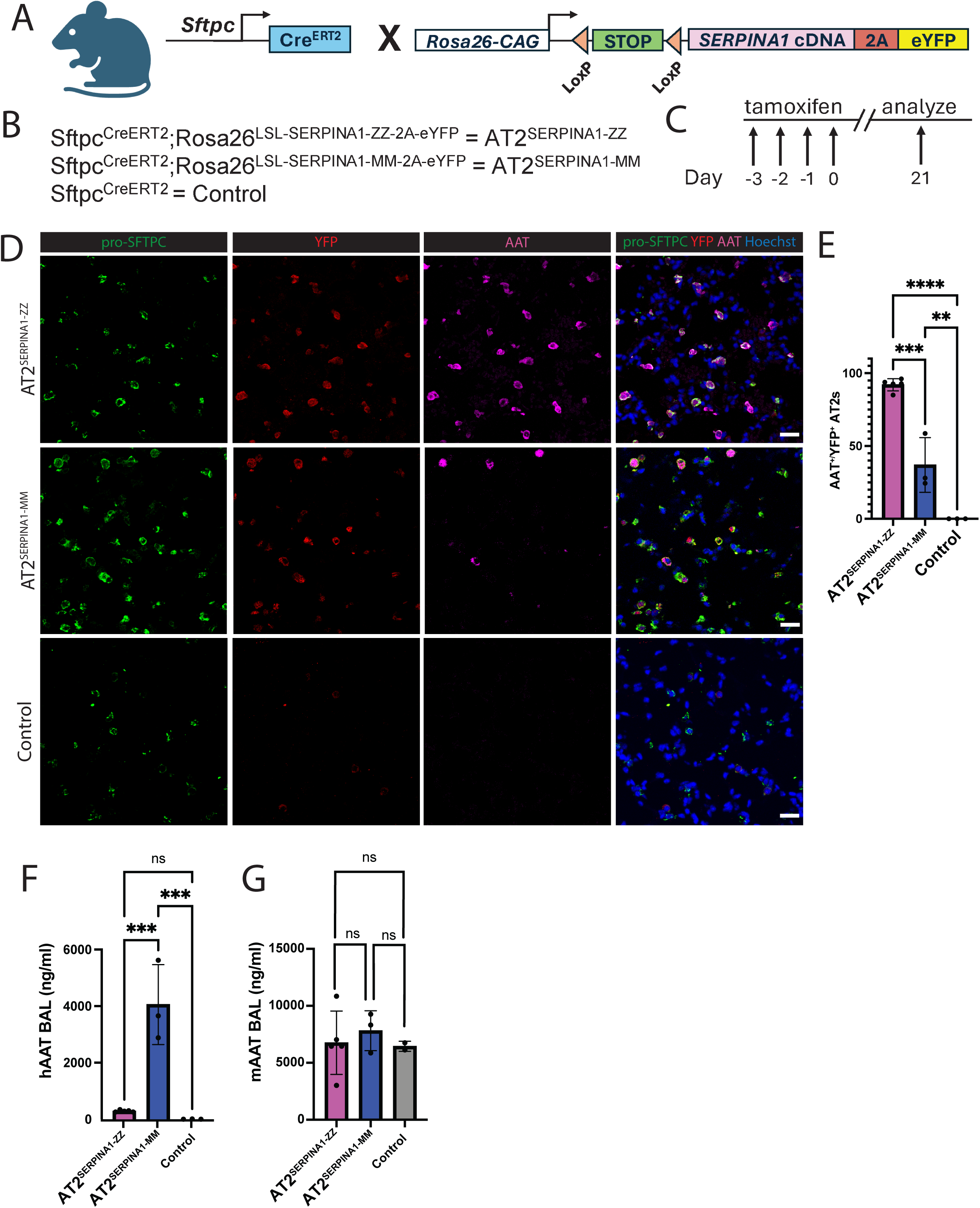
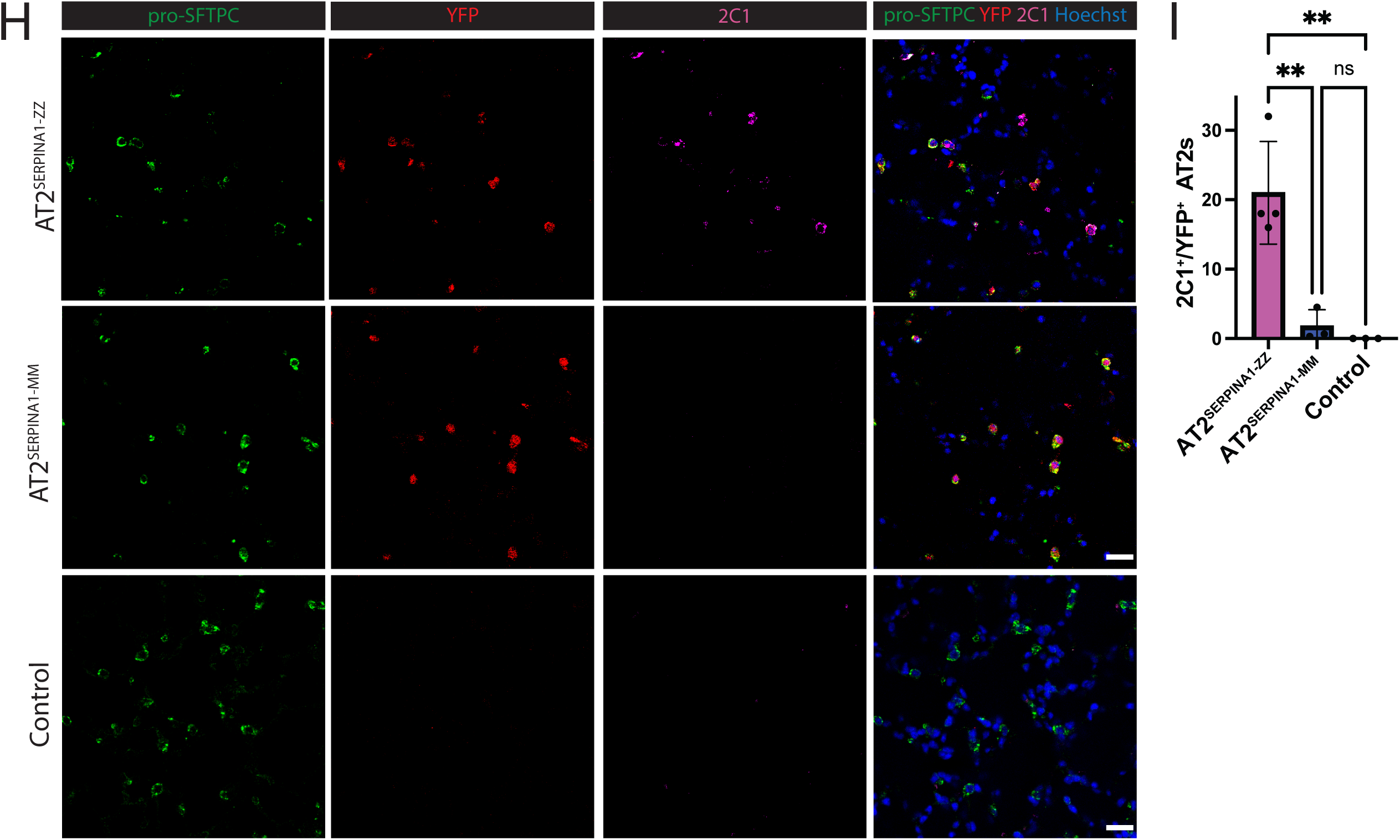
AT2^SERPINA1−ZZ^ mice exhibit increased human intracellular AAT retention and reduced secretion. A) Map of SFTPC^Cre/ERT2^ and Rosa26^LSL-SERPINA1-MM/ZZ-2A-eYFP^ mice crossed to generate experimental mice. B) Mouse nomenclature. C) Experimental design. AT2^SERPINA1-ZZ^, AT2^SERPINA1-MM^, and control mice were injected with tamoxifen once per day over 4 days and then harvested after a 21 day washout. D/H) Immunostaining of AT2^SERPINA1-ZZ^, AT2^SERPINA1-MM^, and control mice for pro-SFTPC, eYFP, and D) AAT or H) 2C1 protein. Scale bar = 25 um. E/I) Mean percent of eYFP+ AT2s staining positive for intracellular E) AAT or I) 2C1 protein in AT2^SERPINA1-ZZ^, AT2^SERPINA1-MM^, and control mice. Each dot represents the mean across multiple fields of view from one mouse. F/G) Secreted total F) human AAT and G) mouse AAT protein in AT2^SERPINA1-ZZ^, AT2^SERPINA1-MM^, and control mouse bronchoalveolar lavage fluid. D-I) AT2^SERPINA1-ZZ^ n=5, AT2^SERPINA1-MM^ n=3, control n=3 and E, F, G, I) Data represented as mean +/- SD. **p < 0.01, ***p < 0.001 by one-way ANOVA with Tukey’s multiple comparisons test.

We first characterized human AAT expression in the lungs of AT2^SERPINA1-ZZ^ and AT2^SERPINA1-MM^ mice. We injected 10 week old mice with tamoxifen over 4 days and harvested lung tissue 3 weeks later for analysis (Figure 3C). Tamoxifen dosing was sufficient to induce eYFP recombination in an average 93% of AT2s (marked by pro-SFTPC), with no evidence of eYFP in control mice, indicating effective recombination in the majority of AT2s (Figure 3D ; Figure S3A,B). Importantly, we only found evidence of eYFP expression in AT2s, defined by expression of pro-SFTPC. We stained tissue sections with an antibody specific for human AAT protein and found that 92% of eYFP+ AT2s expressed human AAT in AT2^SERPINA1-ZZ^ mice compared to 37% in AT2^SERPINA1-MM^ mice and none in control mice, consistent with a reduced efficiency of Z-AAT secretion (Figure 3D,E ; Figure S3A). To further probe the dynamics of AAT retention and secretion, we analyzed levels of human AAT in the bronchoalveolar lavage fluid (BALF) of mice. Consistent with retention of human AAT in AT2^SERPINA1-ZZ^ mice observed by antibody staining, we found lower levels of human AAT in the BALF of AT2^SERPINA1-ZZ^ mice compared to AT2^SERPINA1-MM^ mice (Figure 3F) while murine AAT levels were similar across genotypes (Figure 3G). To determine whether intracellular human AAT forms polymers in mouse AT2s, we next applied the polymer-specific 2C1 monoclonal antibody^36^ and observed polymerized AAT in 21% of eYFP^+^ AT2s from AT2^SERPINA1-ZZ^ mice, compared to only rare staining among eYFP^+^ AT2s from AT2^SERPINA1-MM^ mice (average 2%) and none in control mice (Figure 3H,I ; Figure S3A). These rare 2C1^+^ AT2s in AT2^SERPINA1-MM^ mice might be attributable to high levels of AAT expression, as M-AAT has been shown to polymerize at high concentrations^37^.

Characterization of this novel mouse model demonstrates increased total and polymerized intracellular human AAT in AT2s accompanied by reduced secretion in AT2^SERPINA1-ZZ^ relative to AT2^SERPINA1-MM^ mice. AT2s from AT2^SERPINA1-ZZ^ mice heterogeneously exhibit transcriptomic stress signatures Having observed aberrant AAT processing in AT2^SERPINA1-ZZ^ mouse AT2s, we next wanted to understand the effects of Z-AAT expression on the AT2 transcriptional program. To do so, we injected 10-17 week old AT2^SERPINA1-ZZ^ mice (n=4) and AT2^SERPINA1-MM^ mice (n=3) with tamoxifen. We combined AT2^SERPINA1-ZZ^ mice injected with corn oil (n=3) and Sftpc^Cre/ERT2^ mice injected with tamoxifen (n=3) into a single control group for the purposes of this analysis. Mouse lungs were harvested 8 weeks post-tamoxifen and live cells were sorted, separated into CD45^+^/CD31^+^ immune and endothelial cells, EPCAM^+^ epithelial cells, and triple negative mesenchymal cells, and re-mixed in a 1:1:1 ratio for scRNA-seq (Figure 4A, Figure S4A).

**Figure 4:**
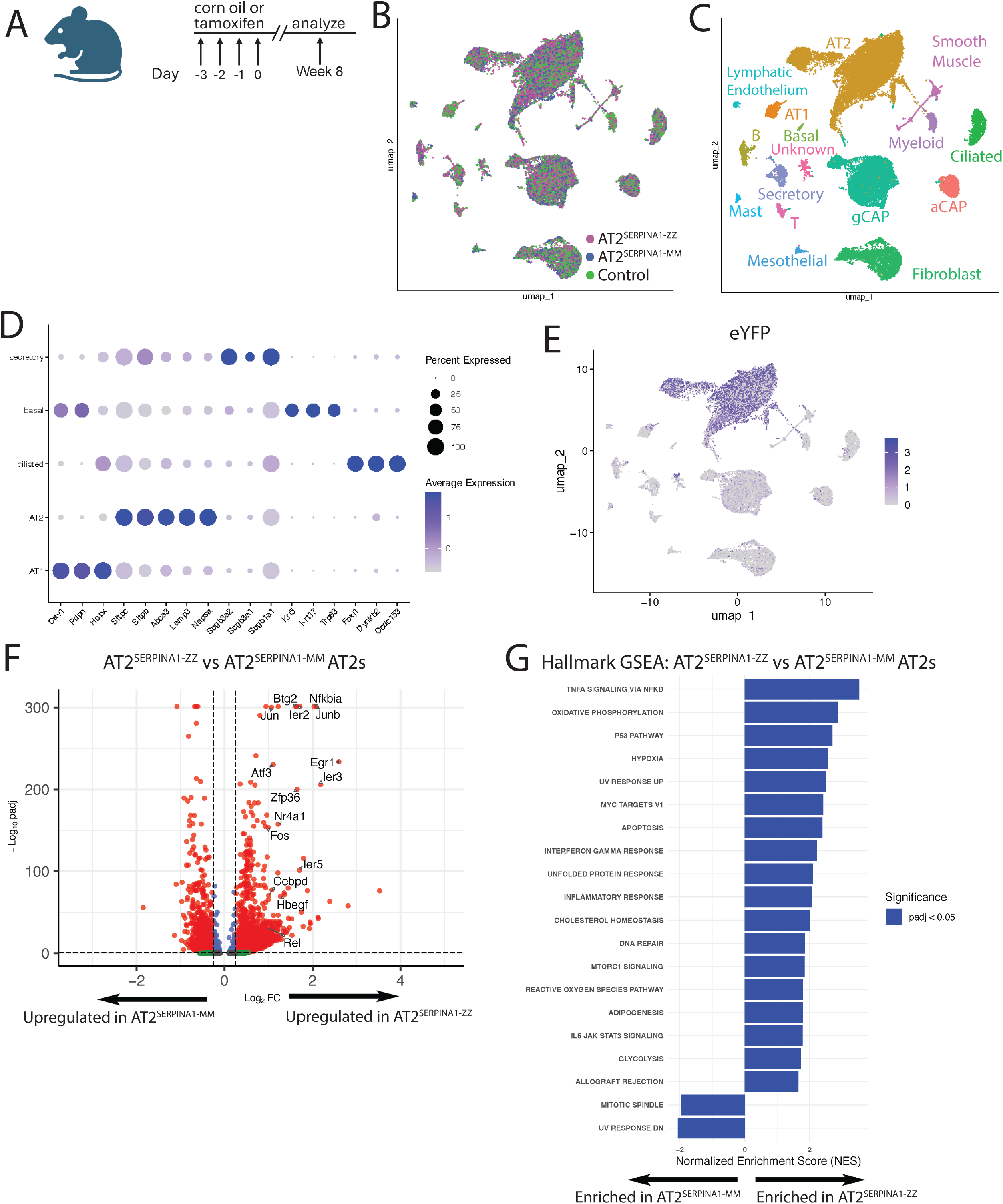
AT2^SERPINA1−ZZ^ mouse AT2s exhibit a transcriptomic disease signature. A) Experimental plan. Mice underwent tamoxifen or corn oil injection followed by scRNA-sequencing of lung tissue 8 weeks later. B) UMAP of AT2^SERPINA1-ZZ^, AT2S^ERPINA1-MM^, and control lung scRNA-seq transcriptomes labeled by genotype. C) UMAP of AT2^SERPINA1-ZZ^, AT2^SERPINA1-MM^, and control lung scRNA-seq transcriptomes labeled by cell identity. D) Dot plot projection for pulmonary epithelial marker gene expression among pulmonary epithelial cell clusters. E) Feature plot of eYFP expression and distribution in UMAP. F) Volcano plot of DEGs comparing AT2^SERPINA1-ZZ^ AT2s versus AT2^SERPINA1-MM^ AT2s by scRNA-seq. Dashed lines show padj < .05 and log2FC > .25. A subset of enriched genes is highlighted with large font. G) Hallmark GSEA of AT2^SERPINA1-ZZ^ AT2s versus AT2^SERPINA1-MM^ AT2s ranked by NES. A-G) AT2^SERPINA1-ZZ^ n=4, AT2^SERPINA1-MM^ n=3, control n=6

Cells clustered primarily based on cell lineage, and not genotype, in UMAP space (Figure 4B,C). We identified multiple epithelial, endothelial, immune, and mesenchymal lineages within each lung (Figure 4C), with epithelial lineages clearly defined by expression of canonical markers (Figure 4D). eYFP expression was mostly constrained to AT2s, and minimally observed in other cell lineages (Figure 4E). Focusing on the AT2 transcriptome, we began by examining differentially expressed genes and identified 1926 upregulated transcripts and 768 downregulated transcripts in AT2^SERPINA1-ZZ^ compared to AT2^SERPINA1-MM^ AT2s. Included in the list of upregulated transcripts were members of the AP-1 complex^38,39^ and stress-responsive genes including *Atf3, Jun, Junb, Fos, Nfkbia, Rel, and Ier3*. Hallmark FGSEA showed enrichment of gene sets including “TNFα signaling via NF-κB”, “Hypoxia”, “Apoptosis”, and “Unfolded Protein Response” (Figure 4F,G). Comparison of AT2^SERPINA1-ZZ^ to control AT2s identified fewer differentially expressed genes with a similar pattern of enrichment, albeit to a lesser degree of significance (Figure S4B,D). Notably, one of the top differentially upregulated genes in this analysis was *Hspa5*, which encodes BiP, the chaperone protein and regulator of ER homeostasis known to be upregulated in human Z-AAT-expressing hepatocytes^40^. Interestingly, comparison of AT2^SERPINA1-MM^ to control AT2s demonstrated reduced expression of cell stress genes and pathways (Figure S4C,E), potentially indicating that expression of M-AAT from AT2s has a protective effect against cell stress.

### Transcriptomic analysis of human AATD AT2s reveals enrichment of proteotoxic stress and inflammatory signatures

We previously published an analysis of lung explant tissue collected from patients at the University of Pennsylvania (UPenn), comparing gene expression patterns in AT2s isolated from PiZZ (“ZZ”) AATD/COPD lungs to those from “MM” COPD or non-diseased controls^13^. To extend these findings in an independent dataset, we reanalyzed data from a recently published dataset of single nucleus RNA sequencing (snRNA-sequencing)^41^ performed on COPD lung tissue samples from the NHLBI Lung Tissue Research Consortium (LTRC). From the 141 LTRC study participants, we identified 5 with severe (ZZ) alpha-1 antitrypsin deficiency based on sequence of *SERPINA1* transcripts. We then analyzed AT2 transcriptional data from these ZZ-COPD samples compared to 15 MM-COPD LTRC samples from former smokers matched on age and FEV to identify differentially expressed genes (Figure 5A). We found 725 genes to be upregulated and 1,324 genes to be downregulated in ZZ-COPD AT2s compared to MM-COPD AT2s (Figure 5B). Among the most upregulated transcripts were *NFKBIZ*, a regulator of NF-κB signaling, the inflammatory chemokine gene *CXCL2*, and the unfolded protein response (UPR)-associated transcription factor *XBP1*. Gene set enrichment analysis (GSEA) identified enrichment of the Hallmark gene set “TNFα Signaling via NF-κB” together with “Hypoxia”, “p53 Pathway” and others (Figure 5C), consistent with our previous findings^13^. Inferred pathway activity analysis via PROGENy similarly showed enrichment of pathways “Hypoxia”, “TNFa”, and “NFkB” in ZZ-COPD AT2s^42^.

**Figure 5:**
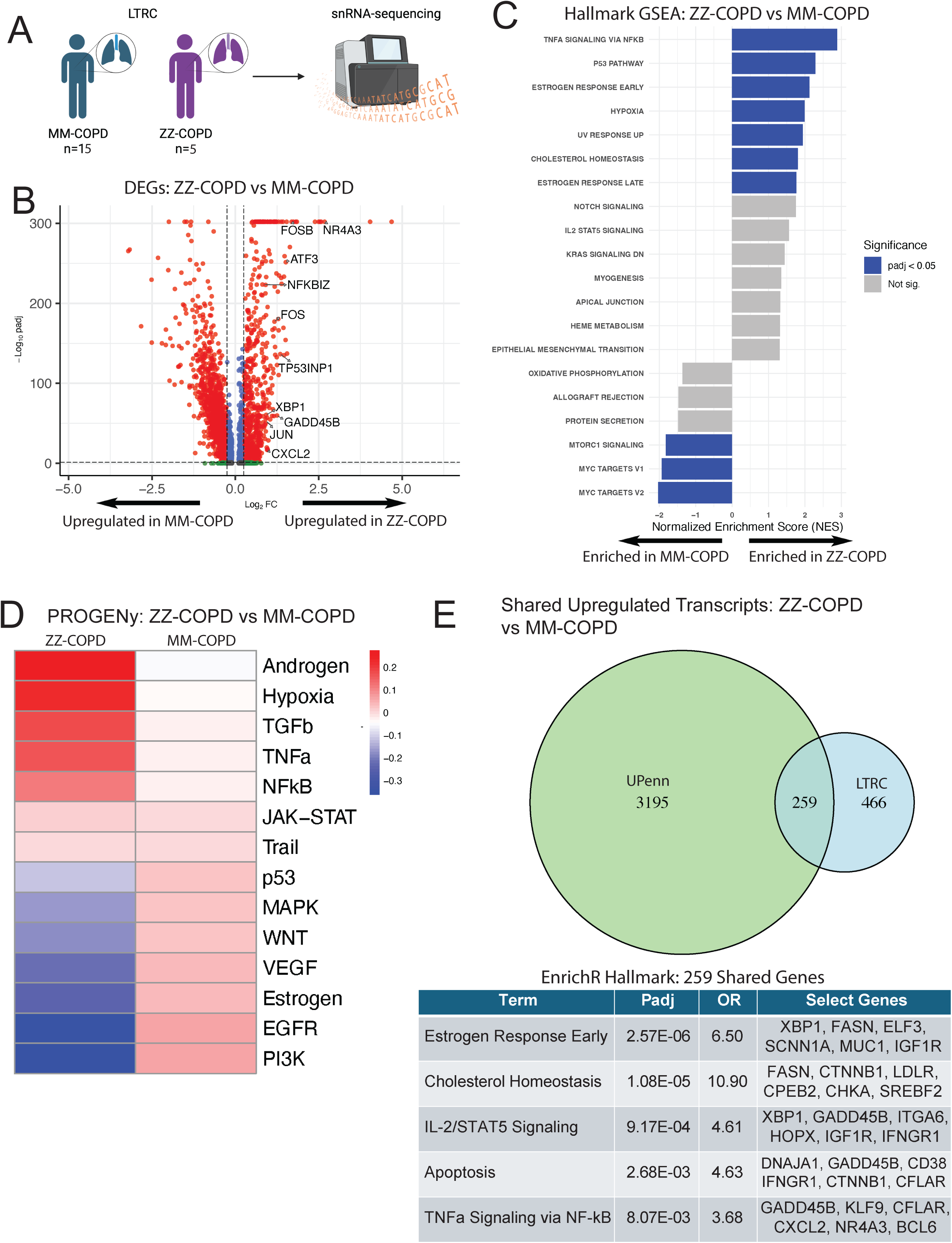
Primary human lung ZZ-COPD AT2s exhibit a transcriptomic disease signature compared to MM-COPD AT2s. A) Experimental outline. Lungs from 15 MM-COPD patients and 5 ZZ-COPD patients from the LTRC underwent snRNA-sequencing. B) Volcano plot of differentially expressed genes (DEGs) comparing LTRC ZZ-COPD AT2s versus wild-type MM-COPD AT2s by scRNA-seq. Dashed lines show padj < .05 and log2FC > .25. A subset of upregulated genes is highlighted with large font. C) Hallmark gene set enrichment analysis (GSEA) of LTRC ZZ-COPD AT2s versus wild-type MM-COPD AT2s ranked by normalized enrichment score (NES). D) PROGENy analysis of LTRC ZZ-COPD AT2s vs MM-COPD AT2s. E) Venn diagram showing upregulated genes (padj < .05, log2FC > .25) comparing UPenn and LTRC ZZ-COPD AT2s versus wild-type MM-COPD AT2s with top Hallmark pathways by EnrichR of the 259 genes enriched in both datasets. A-E) LTRC ZZ-COPD n=5, wild-type MM-COPD n=15 and E) UPenn ZZ-COPD n=2, wild-type MM-COPD n=4.

We next revisited our prior analysis, comparing gene expression in ZZ-COPD to MM-COPD AT2 samples to identify significantly upregulated (n=3454) and downregulated (n=618) genes in ZZ-COPD AT2s (Figure S5A). Cross-referencing this list with differentially expressed genes from the LTRC dataset, we found that 259 genes were upregulated among ZZ-COPD AT2s from both datasets (Figure 5E). Pathway analysis performed on shared upregulated genes revealed enrichment for inflammatory and apoptotic pathways (Hallmark “TNFα Signaling via NF-κB” and “Apoptosis”), consistent with our broader analysis of differential gene expression (Figure 5E).

Specific genes replicated in the two datasets included the apoptosis regulators *CFLAR* and *GADD45B*, chaperone protein encoding gene *DNAJA1,* AP-1 complex gene *JUND,* collagen gene *COL4A4*, and inflammatory cytokine *CXCL2.* Together, these findings are consistent with activation of stress-inflammatory programs in ZZ-COPD AT2s isolated from patients with advanced lung disease (GOLD stage IV) above levels identified in MM-COPD.

### Multiple models of Z-AAT expressing AT2s identify transcriptomic dysregulation of shared pathways

Having generated transcriptional data from human iAT2s, mouse AT2s, and human primary AT2s, we next looked to identify common and distinct genes and pathways that might illuminate disease mechanisms. To do so, we first generated lists of genes significantly upregulated in human primary ZZ-COPD AT2s, ZZ iAT2s, and AT2^SERPINA1-ZZ^ AT2s compared to controls. Comparing expression patterns among these systems, we identified 324 genes upregulated in ZZ iAT2s and AT2^SERPINA1-ZZ^ mice mapping to GO terms such as “Response to Unfolded Protein” and “Regulation of Apoptotic Process” (Figure 6A, Figure S6A). 592 genes were upregulated in AT2s in vivo from human primary ZZ-COPD and AT2^SERPINA1-ZZ^ mice, mapping to GO terms such as “Regulation of apoptotic process” and “Response to endoplasmic reticulum stress” (Figure 6A, Figure S6B). Finally, 905 upregulated genes such as *XBP1*, *JUN, HMOX1,* and *PTEN* were in common between human AT2 systems (primary human ZZ-COPD AT2s and ZZ iAT2s) and mapped to GO terms such as “Regulation of gene expression” (Figure 6A, Figure S6C). Next, we looked across all 3 systems and identified a list of 137 genes that were upregulated in common among Z-AAT expressing AT2s, including multiple genes directly and/or indirectly associated with oxidative stress (*SOD2, HMOX1*), DNA damage (*GADD45A, GADD45B*), AP-1 transcription (*FOS, FOSB, JUN, JUND*), NF-kB signaling (*REL, NFKBIZ, BCL3,* and *ZFAND5*), UPR (*XBP1*), and epithelial remodeling (*KRT8, KRT18*) (Figure 6A). EnrichR analysis of these 137 genes mapped their expression to GO:BP pathways such as “Negative Regulation of Apoptotic Process, “Regulation of Canonical NF-kB Signal Transduction”, and “Positive Regulation of Canonical NF-kB Signal Transduction” (Figure 6B). Evaluation of FGSEA across datasets indicated a larger pattern of pathway enrichment across models, with Hallmark pathways such as “TNFα signaling via NF-κB”, “Hypoxia”, “Apoptosis”, and “Inflammatory Response” either significantly or non-significantly enriched across all model systems (Figure 6C). Application of a previously published gene module for NF-κB signaling^32^ demonstrated a higher module score across models in ZZ compared to MM AT2s, further supporting upregulation of NF-κB signaling as reflected by enrichment of the “TNFα signaling via NF-κB” Hallmark gene set (Figure 6D, Figure S6D). To further investigate this stress-inflammatory phenotype, we applied a gene module describing an “inflammatory AT2” (AT2i) state defined by Zhang et al^41^, across models in ZZ compared to MM AT2s, with the most notable increased expression in UPenn ZZ-COPD AT2s and cluster 4: proteotoxic stress iAT2s (Figure 6E, Figure S6E). To identify drivers of this module score we looked at expression of a select number individual genes composing the module and found them to be broadly upregulated across models (Figure 6F). Together, these data consistently identify NF-κB signaling and ER stress-associated gene expression in Z-AAT-expressing AT2s across model systems, with less consistent dysregulation of other pathways that may reflect context-specificity.

**Figure 6:**
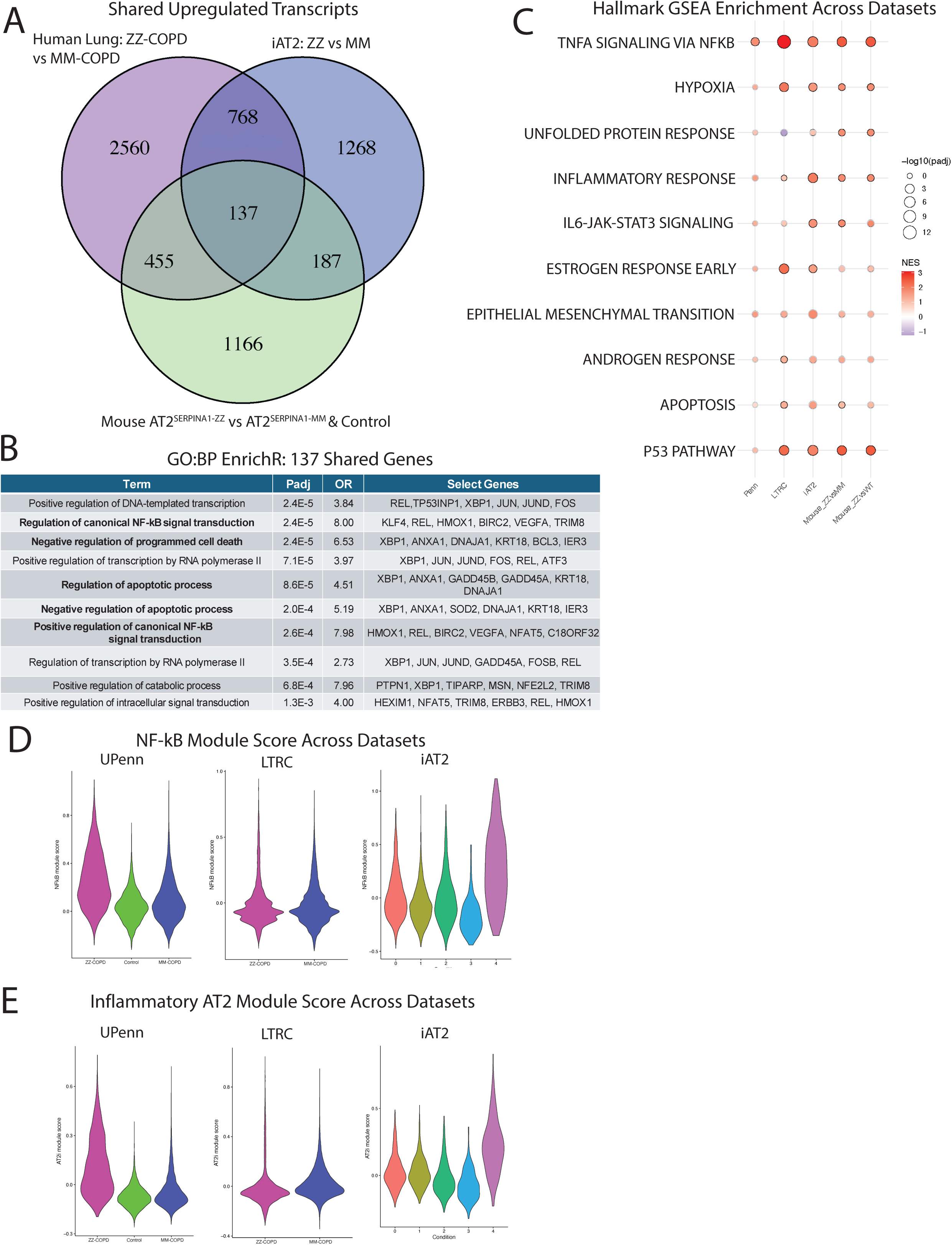
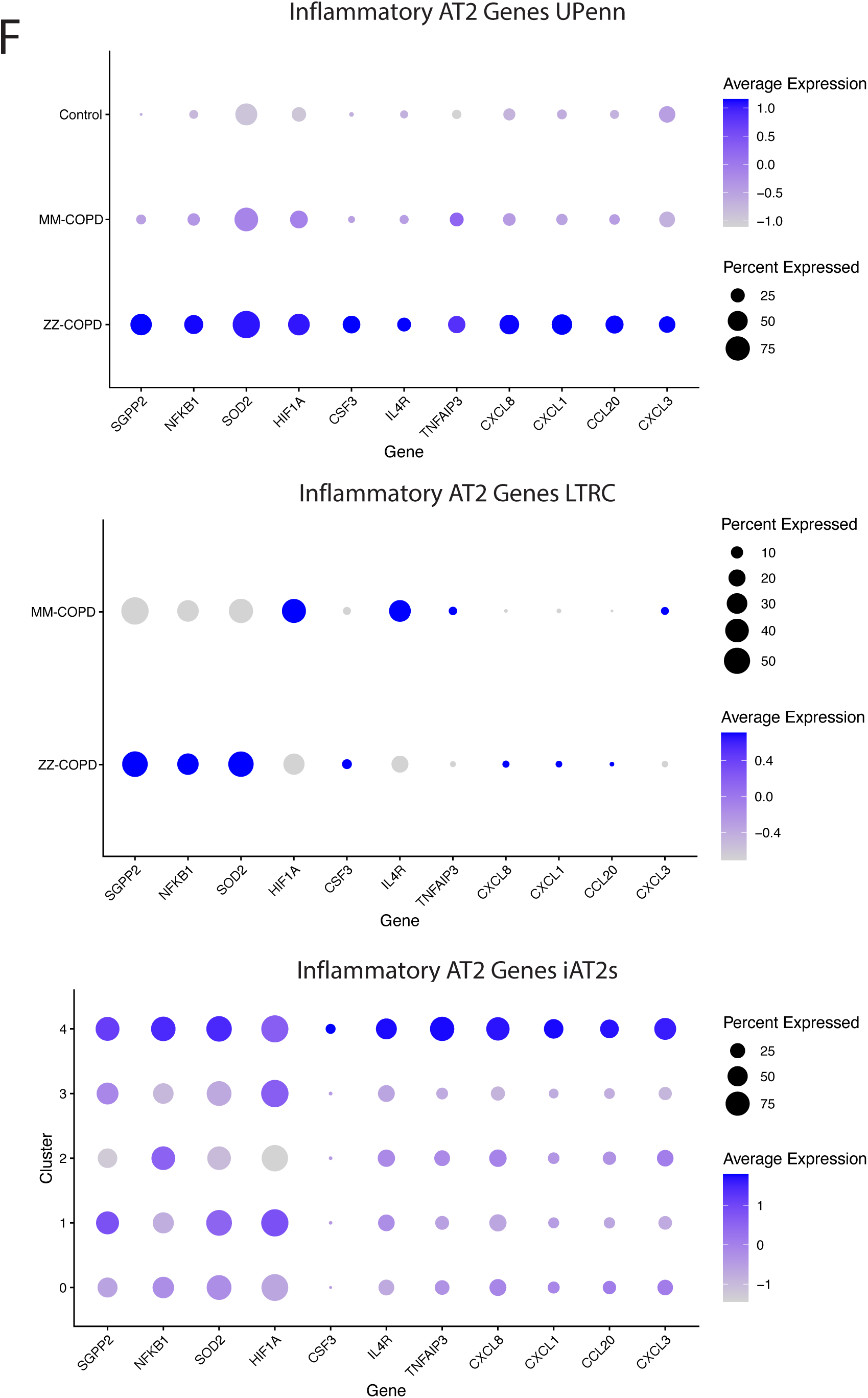
Z-AAT-induced AT2 cellular stress signatures converge across model systems. A) Venn diagram showing upregulated genes (padj < .05, log2FC > .25) comparing human lung AT2s (LTRC and UPenn ZZ-COPD vs MM-COPD), iAT2s (ZZ vs MM), and mouse AT2s (AT2^SERPINA1-ZZ^ versus AT2^SERPINA1-MM^ and AT2^SERPINA1-ZZ^ versus control). B) Top GO:BP terms by EnrichR of the 137 genes upregulated in all datasets. C) Dot plot projection for Hallmark GSEA enrichment across datasets. D) Violin plot of NF- B module score across UPenn, LTRC, and iAT2 datasets. E) Violin plot of inflammatory AT2 module score across UPenn, LTRC, and iAT2 datasets. F) Dot plot projection for inflammatory AT2 module-specific genes across UPenn, LTRC, and iAT2 datasets.

### SERPINA1-ZZ mice are more susceptible to experimental emphysema

We next looked to determine whether the Z-AAT-associated transcriptional disease signature observed in SERPINA1-ZZ mice resulted in a functional impairment with relevance to human AATD-associated lung disease. To do so, we first injured 17-22 week old AT2^SERPINA1-ZZ^ (n=17), AT2^SERPINA1-MM^ (n=5), and control mice (AT2^SERPINA1-ZZ^ mice treated with corn oil (n=7) or mice with the SERPINA1 insert not crossed with Sftpc^Cre/ERT2^ treated with tamoxifen (n=2)) with intratracheal porcine pancreatic elastase (PPE), a well-established model of experimental emphysema^43^ (Figure 7A). Analysis of H&E stained tissue sections demonstrated areas of alveolar enlargement in all three groups that were more pronounced in AT2^SERPINA1-ZZ^ mice compared to other groups (Figure 7B). We quantified mean linear intercept (MLI), a measure of airspace size used to measure emphysema severity, and found increased MLI in AT2^SERPINA1-ZZ^ mice relative to replicate-matched AT2^SERPINA1-MM^ and control mice (Figure 7C). These data indicate Z-AAT expression in murine AT2s increases susceptibility to PPE-induced emphysema.

**Figure 7:**
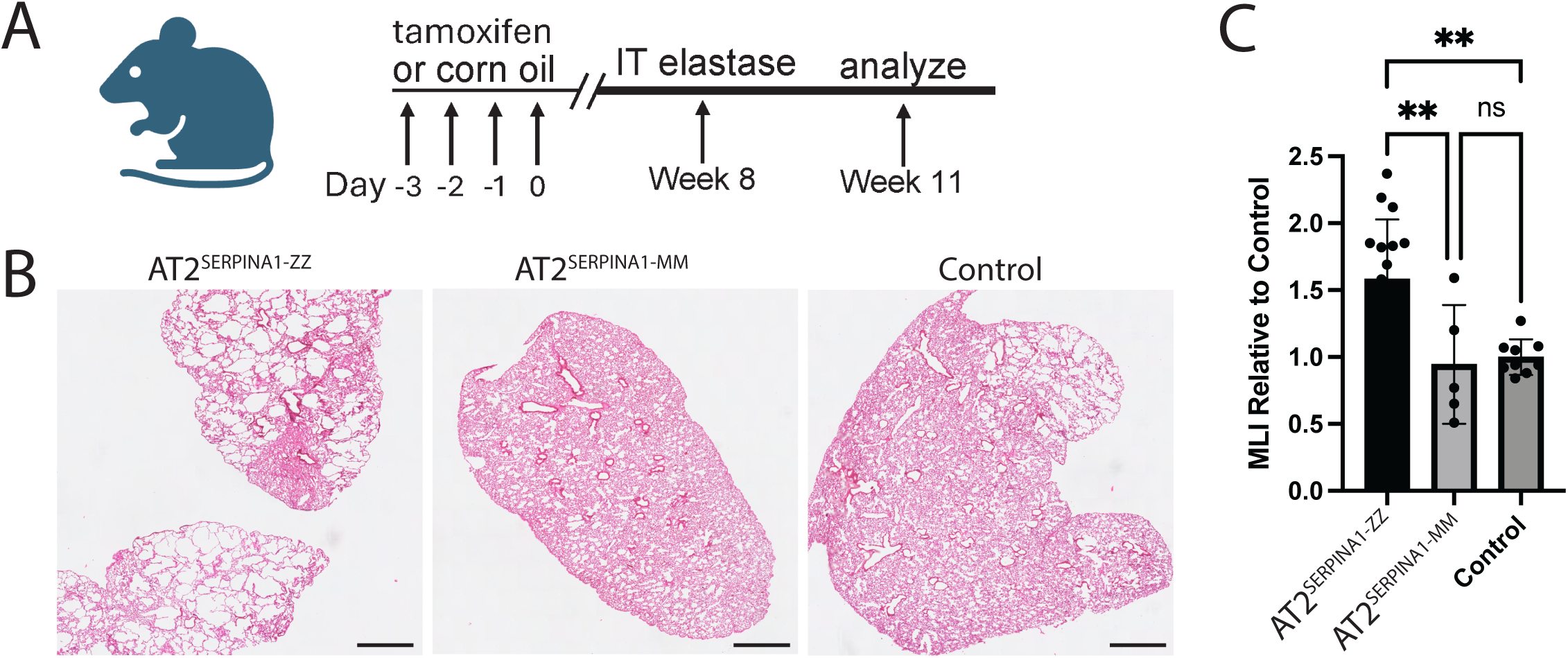
AT2^SERPINA1−ZZ^ mice are more susceptible to elastase-induced emphysema. A) Experimental design. AT2^SERPINA1-ZZ^ and AT2^SERPINA1-MM^ mice were injected with tamoxifen or corn oil once per day over 4 days and then injured with intratracheal (IT) elastase after 8 weeks and harvested 3 weeks after injury. B) Representative H&E images of AT2^SERPINA1-ZZ^, AT2^SERPINA1-MM^, and control mouse lungs 3 weeks after elastase injury. Scale bar = 1 mm. C) Mean linear intercept (MLI) in AT2^SERPINA1-ZZ^, AT2^SERPINA1-MM^, and control mice. Each dot represents the mean across multiple fields of view from one mouse normalized to the mean MLI of control mice from the corresponding technical replicate. n=3 technical replicates. Data represented as mean +/- SD. A-C) AT2^SERPINA1-ZZ^ n=17, AT2^SERPINA1-MM^ n=5, control n=9, p** 0 < .01 by one-way ANOVA with Tukey’s multiple comparisons test.

## Discussion

In this study, we integrated analyses of human iPSC-derived AT2s, a novel conditional expression mouse model, and primary human lung tissue to interrogate the cell-intrinsic consequences of Z-AAT expression in AT2s. Across these complementary model systems, we identified a consistent transcriptional signature characterized by innate immune and inflammatory signaling, NF-κB activation, and ER stress-associated genes and pathways that resulted in increased susceptibility to experimental emphysema in AT2^SERPINA1-ZZ^ mice.

We previously performed the first transcriptional analysis of lung tissue from patients with severe AATD, profiling AT2s from lung explant tissue of two ZZ-COPD donors from a single academic center and identifying dysregulated gene expression when compared to MM-COPD or non-diseased controls. This work now extends that analysis in an independent dataset, utilizing additional patient samples collected at different centers and sequenced using a different platform. We specifically focused on comparisons between ZZ-COPD and MM-COPD to identify molecular features that distinguish lung disease in AATD patients from “usual” COPD. We identified 259 genes that were upregulated in common among Z-AAT expressing AT2s from the two cohorts. Despite technical differences in tissue processing, sequencing approach, and the small sample size that is often a challenge in rare disease research, numerous upregulated gene sets and transcriptional units in the original UPenn dataset were replicated in LTRC samples. GSEA demonstrated enrichment of multiple gene sets in both analyses, including hypoxia, p53 pathway, and TNFα signaling via NF-κB. Notably, these differences emerged despite significantly lower levels of cumulative smoke exposure in the ZZ-COPD participants, highlighting the potential contribution of genetics to this finding.

To determine whether findings observed in primary cells at an advanced disease stage were similarly present in the absence of chronic injury, we applied the iPSC model system, creating syngeneic paired ZZ and MM iPSC lines from four individual donors. In hepatocytes, it is well established that Z-AAT protein “globules” are heterogeneously distributed^44^ and we have similarly observed accumulation of Z-AAT in subsets of AT2s from human tissue^13^. We extend these findings here, noting intracellular AAT staining in a variable proportion of Z-AAT expressing iAT2s, likely a reflection of reduced efficiency of Z-AAT relative to M-AAT secretion. Gene expression patterns in ZZ iAT2s largely recapitulated findings from primary cells, including enrichment for Hallmark gene sets TNFα signaling via NF-κB and inflammatory response.

Further analysis revealed a discrete subcluster of Z-AAT expressing cells characterized by elevated expression of ER chaperone genes, PERK-eIF2α target genes, inflammatory cytokines, and markers of an ABI state, suggesting that proteotoxic stress drives heterogeneous downstream consequences in a subset of Z-AAT expressing AT2s. This heterogeneous response to Z-AAT expression is consistent with observations that we have previously made in Z-AAT expressing iPSC-derived hepatocytes^27^. Taken together, the transcriptional profile of the proteotoxic stress cluster suggests that a subset of Z-AAT expressing AT2s undergo a multifaceted stress response with potential consequences for both inflammatory signaling and cell fate. Interestingly, we did not observe similar transcriptomic heterogeneity within Z-AAT expressing primary human or mouse AT2s, potentially reflecting either the advanced disease stage of human donor lungs or hybrid promoter-driven expression of *SERPINA1* in our mouse model. Although upregulation of PERK effector genes is potentially consistent with activation of the UPR in response to Z-AAT accumulation in the ER, this finding could also represent activation of the integrated stress response (ISR) downstream of oxidative stress and innate immune signaling. Given the concurrent upregulation of these processes in ZZ iAT2s, we cannot distinguish between selective UPR activation and ISR activation without additional protein-level assessment of PERK and eIF2⍺ phosphorylation.

To identify functional consequences of Z-AAT expression in lung epithelium in vivo, we developed a mouse model allowing lineage-specific, inducible human *SERPINA1-*ZZ or *-*MM expression. Unlike prior models, this system induces Z-AAT expression among AT2s. It likewise maintains expression of endogenous murine *Serpina1*, thereby allowing assessment of cell-intrinsic epithelial stress independent of systemic AAT deficiency and the associated reduction in lung antiprotease activity. AT2^SERPINA1-ZZ^ mice demonstrated increased intracellular human AAT retention, reduced secretion into the epithelial lining fluid, and formation of AAT polymers detectable by the polymer-specific 2C1 antibody. These findings recapitulate key features of human Z-AAT biology. Expression of human Z-AAT induced a transcriptional disease signature similar to that observed in human AT2s, both primary and iPSC-derived, suggesting the suitability of this mouse to model features of human disease. While these studies employed Cre recombinase driven from an AT2-specific promoter, future work could apply alternative Cre drivers to explore the proteotoxic effects of Z-AAT expression in other lineages. To directly test whether Z-AAT expression increases susceptibility to lung injury, we injured mice with porcine pancreatic elastase, finding that AT2^SERPINA1-ZZ^ mice developed greater airspace enlargement than controls in response to injury. These findings indicate that even with intact expression of normal, functional AAT to maintain protease-antiprotease balance, epithelial Z-AAT expression is sufficient to increase susceptibility to injury. Further work including the quantification of AT2s following injury will be required to determine whether impaired AT2 regenerative capacity contributes to increased emphysema susceptibility.

Extensive research has previously identified activation of the NF-κB pathway in Z-AAT expressing cells, including primary hepatocytes, monocytes, and neutrophils^1,3,45^. Consistent with these findings, we observed enrichment of the Hallmark gene sets “TNFα signaling via NF-κB”, a previously published module score for NF-κB signaling^32^, and upregulation of the genes *REL* and *NFKBIZ* in Z-AAT expressing mouse and human AT2s and iAT2s. We identified a list of 137 genes that were upregulated in Z-AAT expressing primary human and mouse AT2s and human iAT2s. Included in this list were *SOD2*, an indicator of oxidative stress upregulated in lung epithelium in usual “MM” COPD^41^, *JUND, JUN, FOSB* and *FOS*, AP-1 transcription factors implicated in the epithelial stress response^46^, *XBP1*, an effector in the IRE1 arm of the UPR^47^, *PTPN1*, an enzyme implicated in regulation of PERK signaling and modulation of inflammatory and metabolic cascades^48^, and *ZFAND5*, involved in ubiquitin-proteasome protein degradation and regulation of NF-κB activity^49^. In addition to overlap of specific genes, we likewise noted enrichment across models of genes associated with ER stress- and NF-κB-associated programs, suggesting dysregulation of these pathways as a defining feature of Z-AAT expression in AT2s.

Zhang et al recently identified markers of an inflammatory AT2 cell state in usual MM-COPD including the genes *SGPP2, NFKB1, SOD2, HIF1A, CSF3, IL4R, TNFAIP3, CXCL8, CXCL1, CCL20,* and *CXCL3* and enrichment for gene sets including TNFα signaling via NF-κB, IFN gamma response, Inflammatory response, and UPR and found their proportion to be correlated with GOLD stage and emphysema severity and inversely correlated with DLCO^50^. Further, this inflamed epithelial transcriptional profile was correlated with neutrophil degranulation proteins and inversely correlated with AAT protein in the matched extracellular matrix based on proteomic profiling of decellularized tissue, suggesting a relationship between proteolytic imbalance and the “AT2i” state. Interestingly, multiple genes from this inflammatory AT2 state were upregulated in ZZ AT2s across models. Notably, IFN hallmark gene sets and multiple AT2i markers were enriched in ZZ iAT2s, a system that does not incorporate immune cells, suggesting that Z-AAT proteotoxic stress is sufficient to drive AT2s towards this inflammatory state through an intrinsic mechanism rather than a paracrine response to immune cell activity. Convergence on a shared transcriptional endpoint through a distinct mechanism is consistent with the heightened predisposition to tobacco smoke injury and emphysema observed in ZZ-COPD relative to MM-COPD patients and the possibility that the inflamed AT2 state could represent a common pathway of alveolar epithelial injury caused by either extrinsic loss-of-function or intrinsic gain-of-function mechanisms. Further work including potential pharmacologic or genetic inhibition of PERK signaling is necessary to determine whether and how this inflammatory signaling characterized by NF-κB and IFN, together with markers of the inflammatory AT2i state, are activated by Z-AAT proteotoxic stress.

Intriguingly, the iAT2 proteotoxic stress cluster additionally demonstrated enrichment for markers of an ABI state, including *KRT17* and other genes characteristic of ABI cells identified in the distal lung in pulmonary fibrosis and COPD. In contrast to findings in MM-COPD where inflammatory AT2 and aberrant basaloid populations occupy distinct transcriptional states^50^, in ZZ iAT2s these signatures occurred within the same cluster, raising the possibility that Z-AAT proteotoxic stress drives a more confluent epithelial injury response. Notably, this ABI transcriptional program emerged cell-autonomously in iAT2s in the absence of mesenchymal co-culture, extending prior observations in which this state was activated in iAT2s co-cultured with fibroblasts^33,51^ or expressing the *SFTPC* BRICHOS (C121G) mutation that similarly results in aggregation of misfolded proteins^52^. The emergence of an ABI transcriptional program in the context of a mutation associated with emphysema is notable given that this cell state has been most extensively characterized in pulmonary fibrosis, where AT2s are similarly implicated as primary injury targets and cigarette smoke is similarly a major risk factor. That shared epithelial injury programs can be activated by distinct upstream mechanisms yet produce divergent structural disease outcomes suggests that factors beyond the AT2 transcriptional response itself, potentially including mesenchymal composition, spatial organization, or microenvironmental cues as suggested by Zhang et al^41^, might determine whether alveolar injury results in fibrotic remodeling or parenchymal destruction. The co-occurrence of PERK-eIF2α axis activation and ABI marker expression within the same iAT2 subpopulation raises the possibility that proteotoxic stress may promote aberrant AT2 fate decisions in addition to driving inflammatory signaling. Whether Z-AAT-associated proteotoxic stress contributes to the emergence of aberrant transitional cell states in vivo and whether this phenomenon contributes to impaired alveolar regeneration observed in ZZ-COPD represent important questions for future investigation.

In summary, our findings across primary human, human iPSC, and murine systems support a model in which Z-AAT expression in AT2s induces heterogeneous aberrant AAT retention driving downstream proteotoxic stress, an ABI transcriptional state, and associated NF-κB and inflammatory signaling in a cell-intrinsic manner. Additionally, we observed functional consequences of Z-AAT expression in the form of increased susceptibility to emphysematous injury in mice. Beyond the classical paradigm of loss-of-function toxicity driven by protease-antiprotease imbalance, our data support a potential gain-of-function contribution of Z-AAT driven AT2 transcriptional dysregulation to AATD-associated emphysema pathogenesis. These findings suggest that AT2-intrinsic Z-AAT toxicity and its downstream consequences, including activation of PERK-eIF2α and adoption of an inflammatory AT2 state, represent potential biomarkers of disease activity and/or targets for therapeutic intervention.

### Limitations

This work includes several limitations. First, our studies identify heterogeneity among Z-AAT expressing iAT2s consistent with prior observations made in primary human tissue. In both cases, more work and larger sample sizes are needed to define the mechanisms responsible for this heterogeneity. These studies rely on transcriptional data to infer activation of downstream pathways. Additional studies are needed to confirm this activation at the protein level and to identify the upstream mechanisms driving ER stress and NF-κB activation. Our studies relied on a mouse model co-expressing murine *Serpina1* together with AT2-specific Z-AAT to identify cell-intrinsic phenotypes and associated increased susceptibility to experimental emphysema.

Further work to examine the cell-intrinsic effects of Z-AAT expression in a mouse lacking concomitant *Serpina1* expression would help to extend and clarify these observations.

## Methods

### Sex as a biological variable

Our study examined male and female subjects in all model systems (iAT2, mouse, and primary human lung), and similar findings are reported for both sexes.

### snRNA-seq

The 20 human lung samples used for snRNA-seq experiments in this study were obtained from the National Heart, Lung, and Blood Institute-sponsored LTRC^53^. Deidentified clinical data were obtained from the Biologic Specimen and Data Repository Coordinating Center (BioLINCC).

Human snap-frozen lung tissue samples were processed as described in Sauler et al^54^. Briefly, nuclei were isolated using the Chromium Nuclei Isolation Kit (10x Genomics), stained with DAPI, and counted (Thermo Fisher Scientific Countess II FL). 20,000 nuclei per sample were loaded on Chip M and processed on the Chromium X controller. Libraries were prepared with the Chromium Next GEM Single Cell 3′ HT kit (v3.1). QC was assessed with the Agilent Bioanalyzer High Sensitivity DNA chip. Libraries were sequenced on the NovaSeq System (Illumina) (paired end = 100 bp, targeting 50,000 reads per cell).

### scRNA-seq

The 6 human lung samples used for scRNA sequencing experiments in this study were obtained through an established protocol (PROPEL, University of Pennsylvania). Distal lung tissue was processed into single cells as described previously^55^, except that to isolate primarily distal lung tissue, pleura, airways, and blood vessels were dissected away before processing. Samples were dissociated and CD45-reduced using MACSs LS columns and CD45-microbeads (Miltenyi). Cells were loaded on GemCode (10x Genomics) and were sequenced on the HiSeq2500 or NovaSeq system (Illumina).

### Analysis of human lung RNA-sequencing data

Single cell reads were aligned to the reference genome GRCh38 (human) or GRCm38 (mouse) and pre-processed with Cell Ranger version 8.0.1 (10x Genomics) to obtain the matrix of UMI counts per gene per cell. Count matrices were pre-processed using Seurat version 5.2.17^56^ and filtered to remove dead cells (mitochondrial reads >15% in human lung samples and iAT2s and >10% in mouse, cells with less than 200 genes) and potential doublets (calculated by (100 - (number of cells/1000) / 100)). Human lung donor *SERPINA1* genotype was verified by variant calling from RNA sequencing reads. Data were normalized and scaled using the SCTransform function. Integration of individual human lung and mouse samples were performed using Harmony^57^. Linear dimension reduction was performed using principal component analysis (PCA), and the number of PCA dimensions was evaluated and selected based on assessment of an ElbowPlot. Data were clustered using the Louvain algorithm. UMAP algorithm was used to project the cells onto two dimensional coordinates. Differential gene expression was tested using Wilcoxon rank-sum test. Clusters identified by the Louvain algorithm were annotated based on their DEGs and cell identity was assigned using previously published single-cell atlas expression signatures^35,58,59^. In human and mouse lung datasets, AT2 clusters were subsetted and reanalyzed for differential gene expression. DEGs were visualized using Enhanced Volcano^60^ and heatmaps. Gene set enrichment analysis (GSEA) was performed using FGSEA^61^ for Hallmark gene sets^62^ and ER stress-specific GO gene sets^63,64^ in gene lists ranked by average log2FC. Overrepresentation analysis (ORA) was performed using EnrichR^65–67^ for Hallmark gene sets^62^ and GO Biological Process gene sets^63,64^. Normalized enrichment scores for tests with Benjamini-Hochberg-adjusted false discovery rate < 0.05 were visualized using ggplot^68^. Inferred pathway activity analysis was performed using PROGENy and visualized using pheatmap^42^. Shared upregulated genes were visualized using VennDiagram.

### iPSC reprogramming and maintenance

Peripheral blood mononuclear cells were obtained and reprogrammed into iPSCs according to established protocols. Detailed methods are available for download at https://crem.bu.edu/cores-protocols/. The Institutional Review Board of Boston University approved procurement of PBMCs and reprogramming into iPSCs with written informed consent through protocol H-32506. All iPSC lines used in this study demonstrated a normal karyotype (46XY or 46XX) both before and after gene-editing (Figure S2A). iPSC lines were maintained in feeder-free conditions, on growth factor reduced Matrigel (Corning) in 6-well tissue culture dishes (Corning), in mTeSR Plus media (StemCell Technologies) using ReLeSR (StemCell Technologies) or Gentle Cell Dissociation Reagent (StemCell Technologies) for passaging. iPSCs are maintained in a human iPSC repository and are available upon request at http://stemcellbank.bu.edu.

### Generation of AATD syngeneic iPSC lines

The CRISPR/Cas9 endonuclease system was used to target the *SERPINA1* sequence in close proximity to the Z mutation site within exon 5 using a previously published protocol^27^. Briefly, to achieve scarless editing, two 70bp ssODN repair templates were synthesized (Integrated DNA Technologies) with sequence homology to the *SERPINA1* locus adjacent to the Z mutation and in the complementary orientation with respect to the gRNA sequence. The donor template includes either the wild type SERPINA1 sequence (T to C) or the Z mutation (T) sequence. Both ssODNs contained a silent mutation (G to A) to insert a new ClaI restriction enzyme digest site and facilitate screening for template incorporation. One additional silent mutation (C to T) was included, to reduce subsequent retargeting by Cas9. The resulting M-ssODN sequence was: 5′ TCT AAA AAC ATG GCC CCA GCA GCT TCA GTC CCT TTT TCA TCG ATG GTC AGC ACA GCC TTA TGC ACG GCC T 3’ (used for PiZZ1, PiZZ2, PiZZ6) and the Z-ssODN sequence: 5′ TCT AAA AAC ATG GCC CCA GCA GCT TCA GTC CCT TTT TTA TCG ATG GTC AGC ACA GCC TTA TGC ACG GCC T 3’ (used for BU3). iPSCs were pretreated with 10 μM Y-27632 (Tocris) for 3 hours. Cells were dissociated into single cell suspension with Gentle Cell Dissociation Reagent (StemCell Technologies). 1 × 10^6^ cells were resuspended in 100 μL P3 solution containing Supplement 1 (Lonza) with 5 μl RNP complex (1.5 ul of 100 μM tracrRNA + 1.5 ul of 100 μM crRNA + 2.1 ul of 61 μM Cas9 nuclease V3), 1.2 ul of 100 μM Alt-R Cas9 Electroporation Enhancer, and 1.2 ul of each 100 μM ssODN (Integrated DNA Technologies). The mixture was then nucleofected using the 4-D nucleofector system (Lonza) code CB-150 and replated at a range of densities on growth factor reduced Matrigel coated 6-well plates in mTeSR Plus media containing CloneR 2 (StemCell Technologies). 24 hours later, media was changed to fresh mTeSR Plus containing CloneR 2. iPSCs were left undisturbed for 4-5 days, or until colonies began to expand, then media was changed to mTeSR Plus alone. After approximately 5 more days, emergent colonies were of sufficient size for individual selection, expansion, and screening using the following PCR primers 5’ AGG GAC CAG CTC AAC CCT TC 3’ and 5’ GAG GCA GAC GTG GAG TGA CG 3’. Clones were characterized by Sanger sequencing and next generation sequencing to confirm donor template incorporation (Figure 2A).

### Directed differentiation of iAT2s

We performed iPSC directed differentiation via definitive endoderm into NKX2.1 lung progenitors using previously described methods^21,22^. Briefly, cells maintained in mTeSR Plus media were differentiated into definitive endoderm using the STEMdiff Definitive Endoderm Kit (StemCell Technologies) for 3 days. Cells were then dissociated with Gentle Cell Dissociation Reagent and passaged into 6-well dishes coated with growth factor reduced Matrigel in “DS/SB” anteriorization media, which consists of complete serum-free differentiation media (cSFDM) base as previously described^21,22^ supplemented with 2 μM Dorsomorphin (“DS”, Stemgent) and 10 μM SB431542 (“SB”, Tocris) and 10 μM Y-27632. After 24 hours, media was changed to “DS/SB” without Y-27632. After anteriorization in “DS/SB” media for 72 total hours, cells were cultured in “CBRa” lung progenitor-induction media for 8-9 more days, as previously published^21,22^. “CBRa” media consists of cSFDM base supplemented with 3 μM CHIR99021 (Tocris), 10 ng/mL recombinant human BMP4 (rhBMP4, R&D Systems), and 100 nM retinoic acid (RA, Sigma). On differentiation day 14-15, live cells were sorted on a cell sorter (MoFlo Astrios EQ) to isolate NKX2.1+ lung progenitors based on CPM^+^ gating^22^. Sorted lung progenitors were resuspended in growth factor reduced 3D Matrigel (Corning) at 400 cells/μL. Distalization of cells was performed in “CK+DCI” media, consisting of cSFDM base supplemented with 3 μM CHIR99021, 10 ng/mL rhKGF (CK) and 50 nM dexamethasone (Sigma), 0.1 mM 8-Bromoadenosine 3,5-cyclic monophosphate sodium salt (Sigma) and 0.1 mM 3-Isobutyl-1-methylxanthine (“IBMX”, Sigma) (DCI). Spheres were passaged without further sorting around day 30 of differentiation and submitted to 5 days of CHIR99021 withdrawal followed by one week of CHIR99021 addback. After these 2 weeks, iAT2 cells were sorted again on CPM (MoFlo Astrios EQ) and resuspended in growth factor reduced 3D Matrigel as before. iAT2s were then maintained through serial passaging by plating in 3D Matrigel droplets every 2 weeks with refeeding every other day with “CK+DCI” media, according to our previously published protocol^21,22^. iAT2 culture quality and purity were monitored by flow cytometry for NKX2.1, and resorted on CPM if necessary.

### Flow cytometry and Fluorescence Activated Cell Sorting (FACS)

Preparation of single cell suspensions of 3D Matrigel-embedded iAT2s for flow cytometry and FACS was achieved by incubation with TryplE Express (Thermo Fisher Scientific) for 15–25 minutes at 37°C. Cells were washed with media containing 10% fetal bovine serum (FBS, Thermo Fisher Scientific). Harvested cells were centrifuged at 300 g for 5 minutes at 4°C and resuspended in FACS buffer containing Hank’s Balanced Salt Solution (Thermo Fisher Scientific), 2% FBS and 10 μM Y-27632 (Tocris) and stained with calcein blue AM (Thermo Fisher Scientific) for dead cell exclusion during flow cytometry. Live cells were sorted on a sorter (MoFlo Astrios EQ) at the Boston University Medical Center Flow Cytometry Core Facility.

Immunostaining of iAT2s in single cell suspension for carboxypeptidase M (CPM) was performed as previously described^22^. Briefly, single cell suspensions were stained on ice in FACS buffer using a primary mouse monoclonal antibody against human CPM (1:200, Fujifilm Wako) for 30 minutes and subsequently stained with secondary Alexa Fluor 647 conjugated antibody (1:500, Invitrogen) for 20 minutes. Gating was based on isotype-stained controls.

Immunostaining for NKX2.1 (1:100, Abcam) and AAT (1:100, Santa Cruz) was performed on single cell suspensions of iAT2s fixed for 10 minutes at 37 C in 1.6% PFA (ThermoFisher Scientific). Staining was performed at RT for 30 minutes in permeabilization buffer (eBioscience), and subsequently stained with secondary Alexa Fluor 488 (1:300, Invitrogen) and Alexa Fluor 647 (1:300, Jackson Immunoresearch) conjugated antibodies. Flow cytometry staining was quantified using the Stratedigm S1000EXI and analyzed with FlowJo v10.6.2 (FlowJo, Tree Star Inc). Flow cytometry plots shown represent single cells after forward-scatter/side-scatter gating to remove debris (Figure S2D).

### iAT2 harvest for immunohistochemistry

Whole 3D Matrigel (Corning) droplets of iAT2s were embedded in 2% low-melt agarose (Sigma) and placed on ice for 20 minutes to solidify. The agarose/Matrigel block was fixed in 4% PFA (Thermo Fisher Scientific) at RT for 3 hours, then dehydrated as follows: 2 x 30 minute washes in PBS, 1 x 30 minute wash in 50% ethanol, 2 x 30 minute washes followed by overnight in 70% ethanol, 1 x 1 hour wash in 80% ethanol, 1 x 1 hour wash in 90% ethanol, and 2 x 30 minute washes followed by overnight wash in 100% ethanol, all at 4°C with rocking. Blocks were embedded in paraffin and sectioned with a microtome.

### Immunohistochemistry

Sections underwent rehydration and citric acid-based antigen retrieval followed by blocking/permeabilization with 0.25% Triton X-100 (MilliporeSigma) and 5% normal donkey or goat serum. Samples were then incubated overnight with a combination of the following primary antibodies: NKX2.1 (1:200, Abcam), AAT (1:100, Santa Cruz), pro-SFTPC (7 Hills, 1:500), GFP (Abcam), and 2C1 (1:50, Hycult). Following incubation with primary antibody, samples were washed in PBS with 0.05% Tween 20 (MilliporeSigma) and then incubated with a combination of the following secondary antibodies conjugated to: Alexa Fluor 488 (1:500, Invitrogen), Alexa Fluor 647 (1:500, Jackson Immunoresearch), Alexa Fluor 546 (1:500, Invitrogen) and nuclear Hoechst 33342 (1:500, Invitrogen) for 1 hour. Finally, samples were again washed with PBS containing 0.05% Tween 20 and coverslips were affixed using ProLong Diamond Antifade mountant (Invitrogen). Cells were imaged using the Leica SP5 confocal microscope and images were processed with Fiji software. All quantification was performed via manual counting on 3 randomly selected fields of view each of 3 non-serial sections.

### Preparation of iAT2s for scRNA-seq

Single cell suspensions of iAT2s were sorted for live cells on a sorter (MoFlo Astrios EQ) at the Boston University Medical Center Flow Cytometry Core Facility. scRNA-seq was performed using the GEM-X Single Cell system (10X Genomics) on the NextSeq 2000 machine (Illumina) at the Single Cell Sequencing Core at Boston University Medical Center according to the manufacturer’s instructions (10X Genomics).

### Mice

All animal procedures used to generate this data were approved by the Institutional Animal Care and Use Committee of Boston University. Mice were housed in groups of two to five animals per cage when possible; if singly housed, mice were supplied with extra enrichment. Mice did not undergo any procedures or breeding before tamoxifen administration or intratracheal elastase administration. SFTPC-CreER^T2^ (strain #028054) mice were purchased from Jackson Laboratories^69^. Rosa26^SERPINA1-ZZ/MM-2A-eYFP^ mice were generated by Jackson Laboratories. For all experiments, male and female mice were used, evenly distributed when possible.

### Creation of Rosa26^SERPINA1-ZZ/MM-2A-eYFP^ mouse line

Using CRISPR-based homology-directed repair in mouse embryonic stem cells, Jackson Laboratories inserted a CAG promoter-lox-stop-lox-SERPINA1 cDNA-P2A-eYFP into the Rosa26 locus in the reverse strand at the location Chr6:113067428-113077333 bp on a C57BL/6J background using the CRISPR guides 5’ ACTGGAGTTGCAGATCACGA 3’ and 5’ GCAGATCACGAGGGAAGAGG 3’. Confirmation of correct insertion was confirmed by long-range PCR and sequencing of founders. Detection of *SERPINA1* insert by PCR was performed using the primers 5’ GGT GAA CTT CAA GAT CCG CC 3’ and 5’ CTT GTA CAG CTC GTC CAT GC 3’ at an annealing temperature of 55C with an expected band at 229 bp. Detection of the wild-type allele by PCR was performed using the primers 5’ TCT CCC AAA GTC GCT CTG AG 3’ and 5’ ACT GGA GTT GCA GAT CAC GA 3’ at an annealing temperature of 55 C with an expected band at 196 bp. DNA construct information is available upon request.

### Tamoxifen Administration

Tamoxifen (Sigma-Aldrich) was suspended in corn oil to prepare a stock concentration of 20 mg/mL and incubated at 37 C overnight while rocking to dissolve. Tamoxifen solution was delivered to mice by intraperitoneal (IP) injection of 10 μL per gram body weight at 200 mg/kg every day for 4 days. Mice were weighed daily for 5 days during/following administration and euthanized if their weight loss exceeded 20% of their original body weight.

### Mouse Lung Tissue Harvesting

Mice were euthanized with a lethal dose of CO_2_ followed by cervical dislocation. Perfusion with PBS through the right ventricle was used to clear the lungs of blood. The trachea was cannulated, and BALF was collected through expulsion followed by aspiration of 1 mL of PBS through a syringe. BALF was centrifuged at 400 g for 7 minutes. Supernatant was collected for later analysis and stored at -80 C.

For downstream immunohistochemistry and histology, lungs were then inflation-fixed with 4% PFA (Thermo Fisher Scientific) in PBS at a pressure of 30 cm. Immersion fixation in 4% PFA was continued overnight at 4°C with rocking. Tissue was then dehydrated as follows: 2 x 30 minute washes in PBS, 2 x 30 minute washes in 50% ethanol, overnight wash in 70% ethanol, 1 x 1 hour wash in 80% ethanol, 1 x 1 hour wash in 90% ethanol, and 1 x 1-hour wash followed by overnight wash in 100% ethanol, all at 4°C with rocking. Lung tissue was paraffin embedded and sectioned using a microtome.

### AAT ELISA

Secreted total human AAT was quantified from mouse BALF supernatant as follows. A 96-well high binding plate was coated with 50 ul/well of AAT capture antibody (Genway) at 2 ul/ml diluted in .05 M carbonate-bicarbonate pH 9.6 overnight at RT while rocking. The plate was washed 3x with wash buffer (.05% Tween 20 in PBS, pH 7.4) and blocked with 250 ul/well of blocking buffer (50 mM Tris, .14 M NaCl, 1% BSA, pH 8.0) for 30 minutes. Plate was washed 3x and 100 ul/well of samples diluted in blocking buffer + .05% Tween were incubated on plate for 1 hour. Plate was washed 3x and 100 ul/well of 5.5 ul/ml of detection antibody (R&D) diluted in blocking buffer + .05% Tween was applied and incubated for 1 hour. Plate was washed 3x and 100 ul/well of streptavidin-HRP A (R&D) diluted 1:200 in blocking buffer + .05% Tween was applied and incubated for 20 minutes protected from light. Plate was washed 3x and 100 ul/well of 1:1 TMB peroxidase substrate and Peroxidase substrate solution B (SeraCare) was applied and incubated for 20 minutes protected from light. 100 ul/well of 2N H_2_SO_4_ was applied to each well and read at 450 nm on a microplate reader. Secreted total murine AAT was quantified from mouse BALF using the mouse A1AT ELISA kit (Immunology Consultants Laboratory Inc.) according to manufacturer’s instructions.

### Isolation of murine lung cells for scRNA-seq

After euthanasia and perfusion, lungs were inflated with 1 ml of digestion buffer (9.5 U/ml Elastase (Worthington) + 20 U/ml Collagenase (Gibco) + 5U/ml Dispase (Gibco) + 2 ul/ml DNase I (Sigma)) and 500 ul 1% low melt agarose (Sigma). Parenchyma was minced using a razor blade and incubated with digestion buffer for 45 minutes at 37C while rocking. Tissue was further dissociated through vigorous pipetting and filtered through 70 μM and 40 μM filters. Red blood cells were lysed using red blood cell lysis buffer (Sigma) and single cells were resuspended in FACS buffer. Single cell suspensions of murine lung cells were stained with 1:500 of the following antibodies: CD45-PE (Biolegend), CD31-PE (Biolegend), Epcam-BV421 (BD Horizon) and 1:100 DRAQ7 live/dead (Abcam). Fragments and doublets were excluded and live cells were sorted as follows on a cell sorter (MoFlo Astrios EQ) at the Boston University Medical Center Flow Cytometry Core Facility: PE^+^/BV421^−^ = endothelial and immune, PE^−^/BV421^+^/YFP^+^ = AT2^SERPINA1-ZZ/MM^ AT2s, PE^−^/BV421^+^/YFP^−^ = AT2^SERPINA1-ZZ/MM^ non-AT2 epithelium and all control epithelium, PE^−^/BV421^−^ = mesenchyme (Figure S5A). Cell populations were remixed in a 1:1:1 ratio. scRNA-seq was performed using the Chromium Single Cell 30 system (10X Genomics) and on chip multiplexing (OCM) on the NextSeq 2000 machine (Illumina) at the Single Cell Sequencing Core at Boston University Medical Center according to the manufacturer’s instructions (10X Genomics).

### Emphysema Model

Eight weeks after tamoxifen treatment, animals received intratracheally instilled porcine pancreatic elastase (0.3 U, Worthington) dissolved in 100 μl of PBS. 3 weeks after injury lungs were harvested for histology as described above and stained with hematoxylin and eosin. Digital images of the slides were obtained (Akoya Biosciences Phenoimager HT) and mean linear intercept was quantified using the Measure MLI plugin for Fiji^70^.

### Statistics

Statistical methods relevant to each figure are outlined in each figure legend. In brief, unpaired, 2-tailed Student’s t tests were used to compare quantitative analyses of 2 groups of n = 3 or more samples, and 1-way ANOVAs with Tukey’s multiple comparisons used to compare 3 or more groups. P value threshold for significance was set at p=0.05. Data represented as mean with error bars representing standard deviation. For transcriptomic analyses, the Benjamini-Hochberg false discover rate adjusted p value threshold for significance was set at padj = .05. Specifics about replicates used in each experiment available in figure legends

### Study approval

The Institutional Review Board of Boston University approved procurement of PBMCs and reprogramming into iPSCs with written informed consent received prior to participation through protocol H-32506. All animal procedures used to generate this data were approved by the Institutional Animal Care and Use Committee of Boston University. The 20 human lung samples used for snRNA-seq experiments in this study were obtained from the National Heart, Lung, and Blood Institute-sponsored LTRC^53^. Deidentified clinical data were obtained from the Biologic Specimen and Data Repository Coordinating Center (BioLINCC). The 6 human lung samples used for scRNA sequencing experiments in this study were obtained through an established protocol (PROPEL, University of Pennsylvania).

## Author Contributions

CM, KMA, and AAW conceived of and designed the study. CM, RG, KMA, JEK, MM, MC, MCB, MS, and AAW contributed to data collection. CM, RG, KMA, JEK, PSB, CVM, FW, MC, MCB, MS, and AAW contributed to data analysis. CM and AAW contributed to manuscript preparation.

## Acknowledgements

The authors would like to thank members of the Wilson Lab and Kostas Alysandratos for helpful scientific discussions. We thank Greg Miller, CReM Laboratory Manager; Marianne James, CReM iPSC Core Manager; and Meenakshi Lakshminarayanan, CReM Administrative Director, for their invaluable support. For technical support, we would like to thank Brian R. Tilton of the Boston University Flow Cytometry Core Facility; Yuriy Aleksyeyev of the Sequencing Core Facility, supported by NIH (1UL1TR001430); Lynn Deng and Matthew Au of the Analytical instrument Core; and Michael T. Kirber of the Cellular Imaging Core. CM would like to thank members of her dissertation advisory committee, Joseph Mizgerd, Finn Hawkins, Xaralabos Varelas, and Katrina Traber. Figure schematics and cartoons were created with BioRender.com. Finally, we thank the alpha-1 community as without their selfless participation, none of this work could be done.

