## Supplemental Figures 1-6 for "AT2-intrinsic Z-AAT expression drives conserved inflammatory and proteotoxic stress responses and predisposes to emphysema"

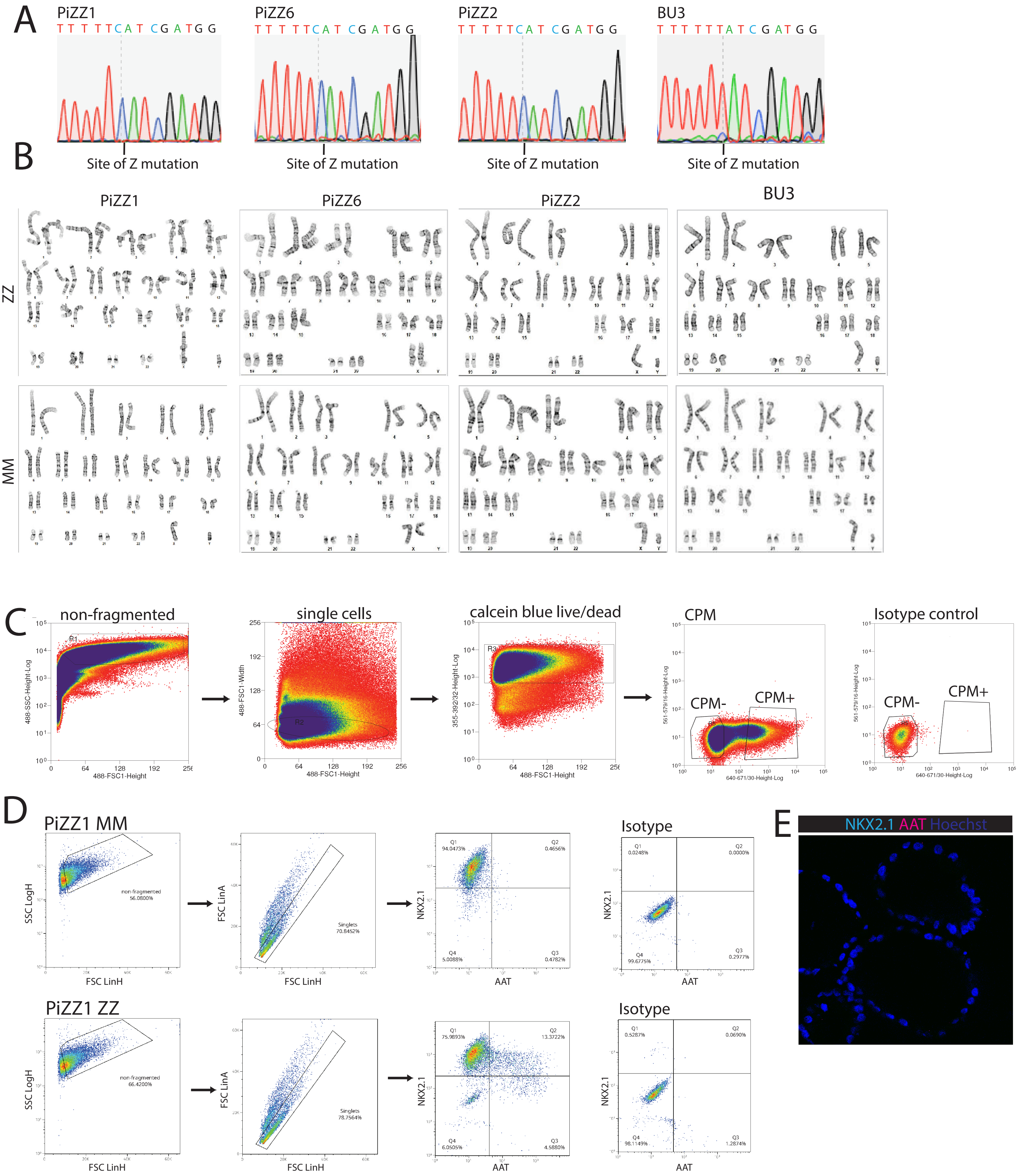
A
PiZZ1
PiZZ6
PiZZ2
BU3
Site of Z mutation
B
PiZZ1
PiZZ6
PiZZ2
BU3
ZZ
MM
C
non-fragmented
single cells
calcein blue live/dead
CPM
Isotype control
CPM-
CPM+
488-SSC-Height-Log
488-FSC1-Height
488-FSC1-Width
355-392/32-Height-Log
561-579/16-Height-Log
640-671/30-Height-Log
D
PiZZ1 MM
PiZZ1 ZZ
Isotype
SSC LogH
FSC LinH
FSC LinA
NKX2.1
AAT
non-fragmented
56.0800%
66.4200%
Singlets
70.8452%
78.7564%
Q1
94.0473%
Q2
0.4656%
Q3
0.4782%
Q4
5.0088%
Q1
0.0248%
Q2
0.0000%
Q3
0.2977%
Q4
99.6775%
Q1
75.9893%
Q2
13.3722%
Q3
4.5880%
Q4
6.0505%
Q1
0.5287%
Q2
0.0690%
Q3
1.2874%
Q4
98.1149%
E
NKX2.1 AAT Hoechst

**Figure S1**

A) Sanger sequencing of iPSC lines after CRISPR-editing.
B) Karyotypes of iPSC lines before and after CRISPR-editing.
C) Representative flow plots and gating of d14 CPM sort in iAT2 differentiation.
D) Representative flow plots and gating for NKX2.1 and AAT staining and isotype controls.
E) Isotype control immunofluorescence staining for NKX2.1 and AAT protein.

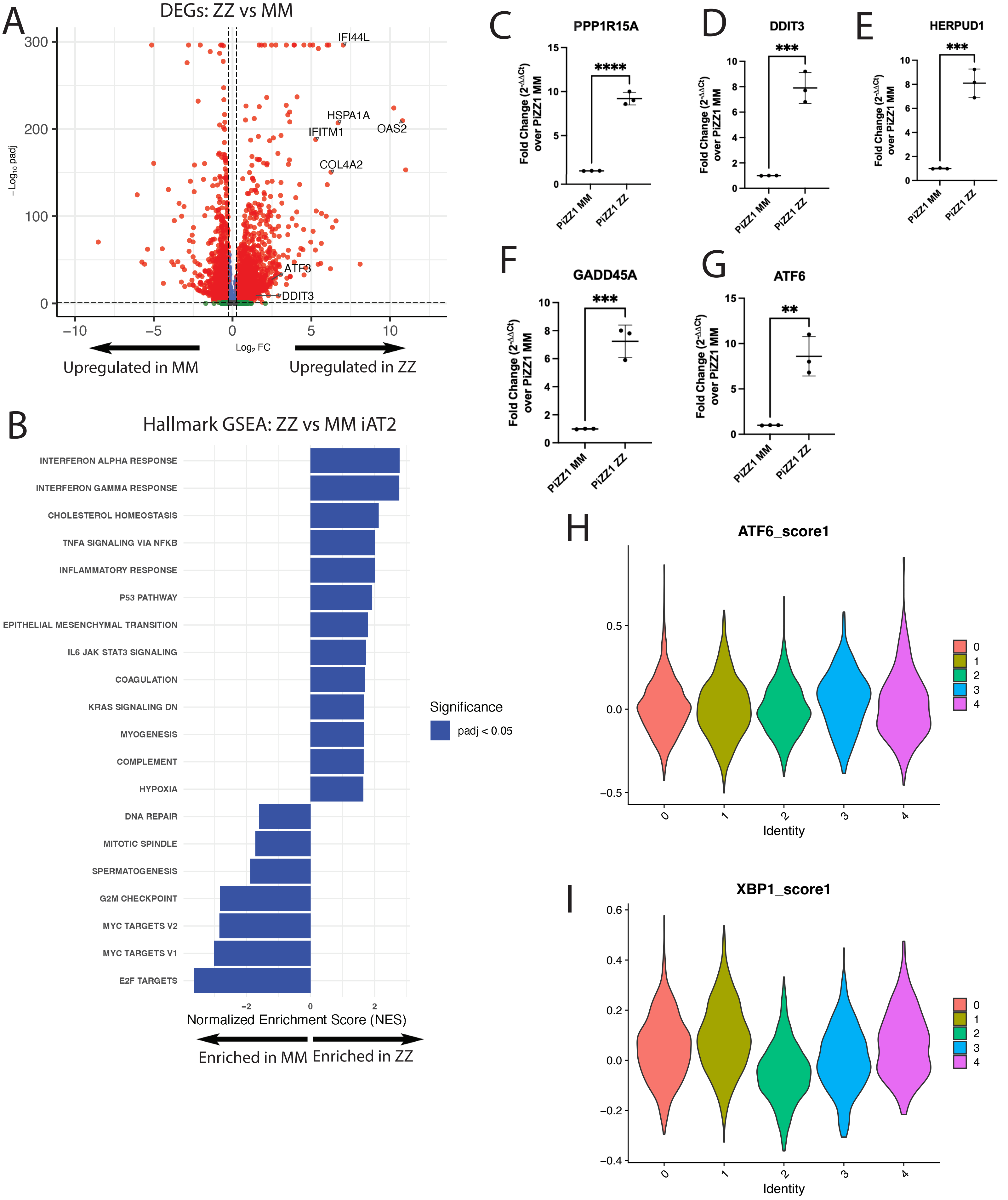
A
DEGs: ZZ vs MM
IFI44L
HSPA1A
OAS2
IFITM1
COL4A2
ATF3
DDIT3
−Log10 padj
Log2 FC
Upregulated in MM
Upregulated in ZZ
B
Hallmark GSEA: ZZ vs MM iAT2
INTERFERON ALPHA RESPONSE
INTERFERON GAMMA RESPONSE
CHOLESTEROL HOMEOSTASIS
TNFA SIGNALING VIA NFKB
INFLAMMATORY RESPONSE
P53 PATHWAY
EPITHELIAL MESENCHYMAL TRANSITION
IL6 JAK STAT3 SIGNALING
COAGULATION
KRAS SIGNALING DN
MYOGENESIS
COMPLEMENT
HYPOXIA
DNA REPAIR
MITOTIC SPINDLE
SPERMATOGENESIS
G2M CHECKPOINT
MYC TARGETS V2
MYC TARGETS V1
E2F TARGETS
Significance
padj < 0.05
Normalized Enrichment Score (NES)
Enriched in MM
Enriched in ZZ
C
PPP1R15A
D
DDIT3
E
HERPUD1
F
GADD45A
G
ATF6
Fold Change (2^-ΔΔCt) over PiZZ1 MM
PiZZ1 MM
PiZZ1 ZZ
H
ATF6_score1
I
XBP1_score1
Identity

**Figure S2**

A)Volcano plot of DEGs comparing ZZ versus MM iAT2s by scRNA-seq. Dashed lines show padj < .05 and log2FC > .25. A subset of upregulated genes is highlighted with large font.
B)Hallmark GSEA of ZZ versus MM iAT2s ranked by normalized enrichment score (NES).
C-G) qRT-PCR of select genes upregulated in scRNA-seq from a separate differentiation. PiZZ1 line, n=3 technical replicates. Data represented as mean +/- SD. **p < 0.01, ***p < 0.001, ****p < 0.0001 by Student's T test.
H) Violin plot of ATF6 module score across all iAT2 clusters.
I) Violin plot of XBP1 module score across all iAT2 clusters.

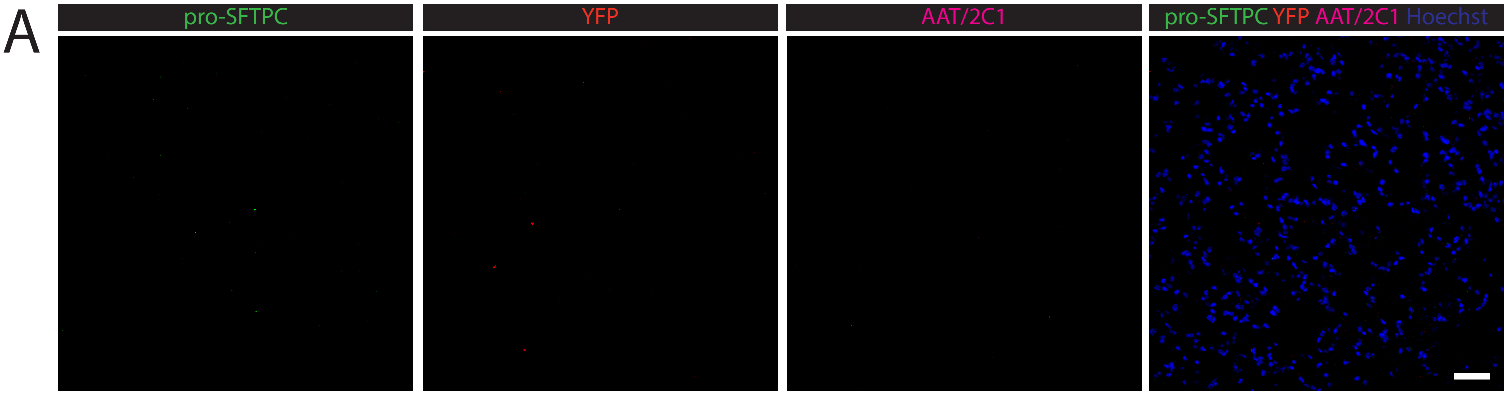


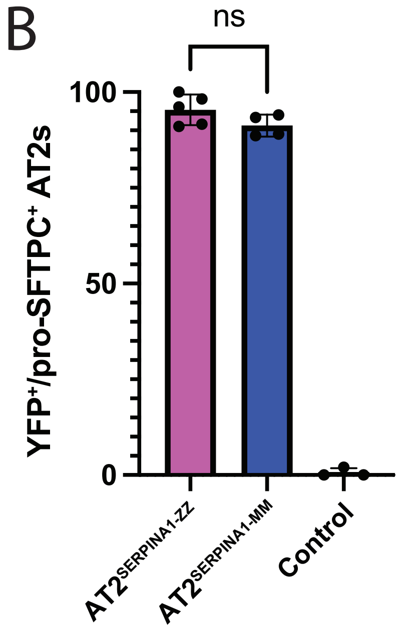


**Figure S3**
A) Isotype control immunofluorescence staining for pro-SFTPC, eYFP, and AAT/2C1 protein.
B) Mean percent of pro-SFTPC+ iAT2s staining positive for eYFP in AT2SERPINA1-ZZ, AT2SERPINA1-MM, and control mice. Each dot represents the mean across multiple fields of view from one mouse. Data represented as mean +/- SD.

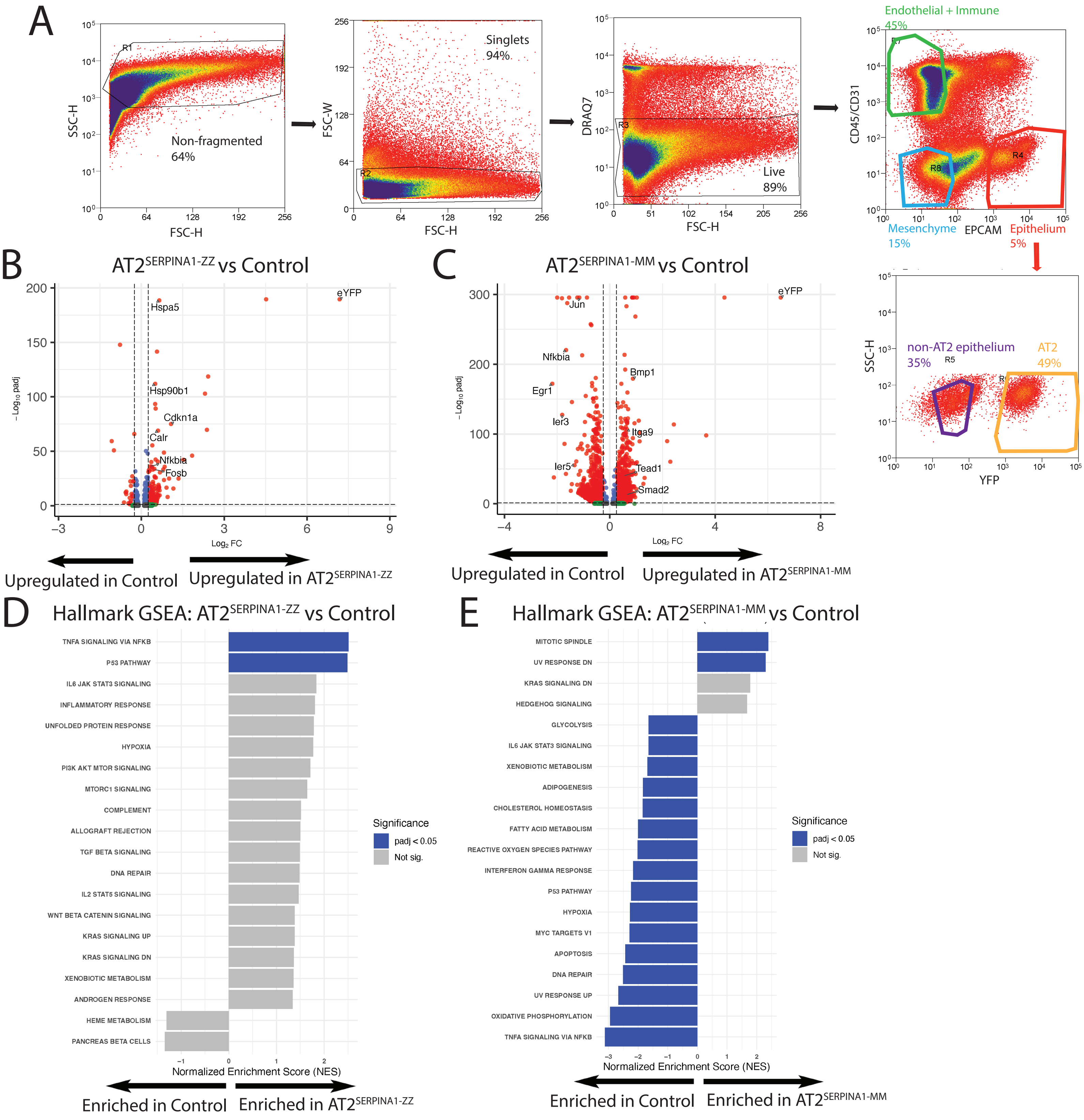


**Figure S4**

A) Representative sort plots for isolating live endothelial, immune, mesenchymal, epithelial, and eYFP+ from mouse lung for scRNA-seq.

B/C) Volcano plot of DEGs comparing B) AT2SERPINA1-ZZ AT2s versus control AT2s and C) AT2SERPINA1-MM AT2s versus control AT2s by scRNA-seq. Dashed lines show padj < .05 and log2FC > .25. A subset of enriched genes is highlighted with large font.

D/E) Hallmark GSEA of D) AT2SERPINA1-ZZ AT2s versus control AT2s and E) AT2SERPINA1-MM AT2s versus control AT2s ranked by normalized enrichment score (NES).

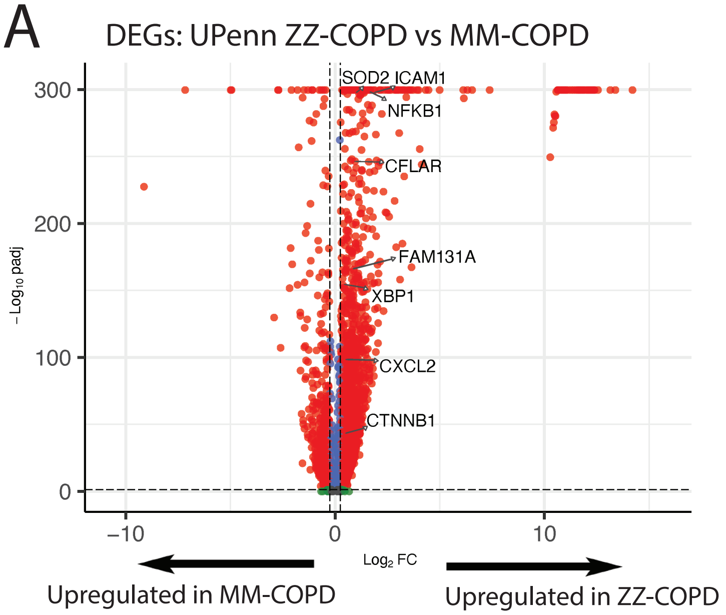


**Figure S5**

A) Volcano plot of differentially expressed genes (DEGs) comparing UPenn ZZ-COPD AT2s versus wild-type MM-COPD AT2s by scRNA-seq. Dashed lines show padj < .05 and log2FC > .25. A subset of upregulated genes is highlighted with large font.

**A**

GO:BP EnrichR: Shared Genes iAT2s and Mouse (324 genes)

| Term | Padj | OR | Select Genes |
|---|---|---|---|
| **Negative regulation of programmed cell death** | 2.0E-5 | 4.32 | XBP1, ANXA1, DNAJA1, KRT18, HMOX1, IER3 |
| **Response to endoplasmic reticulum stress** | 2.0E-05 | 6.03 | PPP1R15A, JUN, XBP1, HERPUD1, DDIT3, ATF4 |
| **Response to unfolded protein** | 2.0E-05 | 16.03 | DNAJA1, DDIT3, HERPUD1, PTPN1, DNAJB4, DNAJB2 |
| **Regulation of apoptotic process** | 2.5E-05 | 3.21 | LGALS3, GADD45A, ATF4, XBP1, JUN, ANXA1 |
| **Negative regulation of apoptotic process** | 1.1E-04 | 3.57 | IER3, XBP1, ANXA1, SOD2, DNAJA1, NUPR1 |
| **Integrated stress response signaling** | 3.5E-04 | 16.07 | CEBPB, JUN, CEBPD, DDIT3, FOS, ATF4 |
| Positive regulation of catabolic process | 5.3E-04 | 5.21 | PTPN1, HERPUD1, CDKN1B, MSN, XBP1, TRIM8 |
| **Positive regulation of autophagy** | 1.2E-03 | 7.01 | TRIM8, CDKN1B, HERPUD1, NFE2L2, XBP1, BNIP3 |
| Negative regulation of DNA-templated transcription | 1.7E-03 | 2.41 | CEBPB, CDKN1B, XBP1, JUN, GADD45A, ATF4 |
| Positive regulation of intracellular signal transduction | 1.7E-03 | 2.77 | PPP1R15A, TRIM8, HMOX1, XBP1, DDIT3, REL |

**B**

GO:BP EnrichR: Shared Genes Primary AT2s and Mouse (592 genes)

| Term | Padj | OR | Select Genes |
|---|---|---|---|
| Positive regulation of DNA-templated transcription | 3.5E-13 | 2.92 | IER2, MAP2K3, JUN, XBP1, JUND, NFKBIA |
| Positive regulation of transcription by RNA polymerase II | 6.1E-12 | 3.05 | AREG, IER2, RELA, JUN, REL, FOS |
| **Regulation of apoptotic process** | 5.1E-10 | 3.22 | IER3, ANXA1, TNFRSF12A, RELA, HSPA5, CALR |
| **Negative regulation of apoptotic process** | 4.8E-7 | 3.25 | ICAM1, SOD2, NFKB1, DNAJA1, KRT18, BIRC3 |
| **Negative regulation of programmed cell death** | 4.8E-7 | 3.57 | HMOX1, IER3, XBP1, TP53, RELA, BIRC3 |
| Negative regulation of DNA-templated transcription | 4.8E-7 | 2.47 | NFKB1, SMAD7, CALR, JUN, JUND, ATF3 |
| Regulation of transcription by RNA polymerase II | 4.9E-7 | 1.97 | HMOX1, GADD45A, REL, CALR, EIF4A1, JUN |
| Negative regulation of transcription by RNA polymerase II | 6.5E-7 | 2.71 | AREG, TP53, ATF3, JUN, XBP1, GADD45A |
| **Response to endoplasmic reticulum stress** | 1.3E-6 | 4.73 | PTPN1, JUN, XBP1, HSPA5, CALR, ATF6 |
| **Regulation of protein ubiquitination** | 1.6E-6 | 6.38 | HSP90AB1, TNFAIP3, DNAJB2, DNAJA1, BIRC2, BIRC3 |

**C**

GO:BP EnrichR: Shared Genes Primary AT2s and iAT2s (905 genes)

| Term | Padj | OR | Select Genes |
|---|---|---|---|
| Positive regulation of DNA-templated transcription | 7.9E-13 | 2.48 | ATF3, JUN, XBP1, KLF6, JUND, JUP |
| Positive regulation of transcription by RNA polymerase II | 7.2E-11 | 2.53 | HIF1A, JUND, JUP, VEGFA, LIF, XBP1 |
| Regulation of transcription by RNA polymerase II | 2.5E-8 | 1.86 | HSPA1A, KLF6, VEGFA, HMOX1, JUN |
| Regulation of DNA-templated transcription | 5.9E-6 | 1.74 | HIF1A, JUND, NCOA2, HMOX1, KLF6, BCL3 |
| Negative regulation of DNA-templated transcription | 5.9E-6 | 2.07 | CTNND1, GADD45A, HSPA1A, HIF1A, XBP1, JUN |
| Regulation of gene expression | 1.3E-5 | 1.97 | NFKBIZ, REL, HSPA1A, HSPA1B, CCNL2, HIF1A |
| Negative regulation of cell motility | 3.2E-5 | 4.27 | SERPINB1, PTEN, DUSP10, CYP1B1, ARID4A, FOXO3 |
| Regulation of cell migration | 3.7E-5 | 2.48 | ARID4B, DUSP10, CASP8, CYP1B1, SERPINB1, ANXA1 |
| Negative regulation of transcription by RNA polymerase II | 6.0E-5 | 2.14 | CTNND1, ATF3, JUN, XBP1, KLF4, FOSB |
| Negative regulation of cell migration | 3.9E-4 | 3.38 | PTEN, ARID4A, ARID4B, DUSP10, SERPINB1, HMOX1 |

**D** NF-kB Module Score Mouse

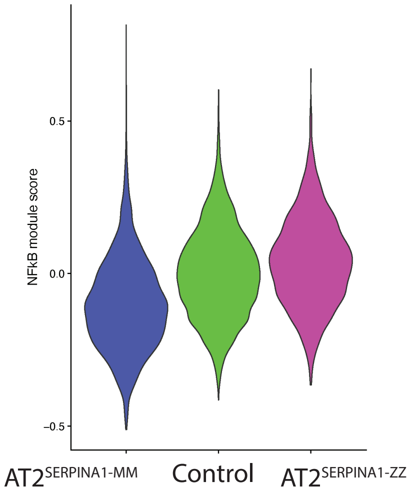


**E** Inflammatory AT2 Module Score Mouse

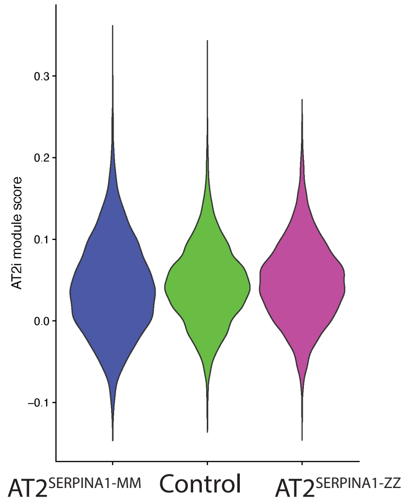

**Figure S6**

A) Top GO:BP terms by EnrichR of the 324  genes upregulated in iAT2 and mouse datasets.

B) Top GO:BP terms by EnrichR of the 592 genes upregulated in human lung and mouse datasets.

C) Top GO:BP terms by EnrichR of the 905 genes upregulated in human lung and iAT2 datasets.

D) Violin plot of NF-  B module score across mouse genotypes.

E) Violin plot of inflammatory AT2 module score across mouse genotypes
